# Deep ancestral structure, admixture and peripheral persistence shape diversity in the *Papilio bianor* species complex

**DOI:** 10.64898/2026.09.21.752948

**Authors:** Haolin Wu, Tonghang Wang, Bo Xu, Youhua Chen

**Affiliations:** Mountain Ecological Restoration and Biodiversity Conservation Key Laboratory of Sichuan Province, Chengdu Institute of Biology, Chinese Academy of Sciences, Chengdu 610213, China; China Croatia Belt and Road Joint Laboratory on Biodiversity and Ecosystem Services, Chengdu Institute of Biology, Chinese Academy of Sciences, Chengdu 610213, China; Department of Biological Sciences, National University of Singapore, Singapore 119077, Singapore; Key Laboratory of National Forestry and Grassland Administration on Biodiversity Conservation on the Qinghai-Xizang Plateau, Chengdu Institute of Biology, Chinese Academy of Sciences, Chengdu 610213, China

## Abstract

Rapid diversification can generate regional biodiversity through ancestral polymorphism, admixture, and ecological differentiation, while leaving complex genomic signals. We analysed whole-genome variation in 325 butterflies, including 166 individuals of the *Papilio bianor* species complex and six outgroup species, to reconstruct lineage relationships and demographic history. Nuclear genomic analyses recovered three major clades corresponding to the recognized species and placed the island lineage as sister to continental *P. bianor*. Analyses of 18,635 autosomal window trees revealed extensive genealogical discordance, with incomplete lineage sorting predominating among continental lineages and the strongest introgression signals involving island-related ancestry. Demographic modelling supported early island–continental divergence followed by continental diversification and ancient admixture. The best-fitting fastsimcoal2 model inferred an admixed origin of the Eastern China lineage, while BPP analyses supported ancient island-related introgression into the Eastern China–Taiwan ancestor, Western Himalaya and southwestern Thailand. Peripheral populations exhibited pronounced genetic differentiation and distinct demographic histories; Western Himalaya additionally showed mitochondrial affinity with Yaeyama despite its placement within the Himalayan nuclear lineage. Together, these findings suggest that introgression among deeply differentiated ancestral populations and incomplete lineage sorting during rapid divergence jointly shaped lineage diversity in the *P. bianor* complex. This study provides an empirical framework for investigating how lineage diversity emerges rapidly during the early stages of speciation, particularly in groups with deep ancestral population structure.

## Introduction

Species diversification has traditionally been represented as a sequence of bifurcations, and in many biological systems population genetic structure broadly corresponds to geographical distribution. Geological barriers and climatic oscillations can restrict dispersal, promote population isolation and ultimately generate genetic, phenotypic and taxonomic diversity^1^. However, evolutionary histories are not always exclusively tree-like. Introgressive hybridization can transfer alleles between previously separated lineages, allopolyploidization can combine complete parental genomes, and horizontal gene transfer can introduce genetic material across distantly related lineages^2,3^. These processes can generate novel genetic backgrounds and phenotypes and, in some cases, contribute directly to adaptation, evolutionary radiation or the formation of reproductively independent lineages. Reticulate evolution should therefore be regarded as an important force, alongside geographical isolation and divergent selection, in shaping biodiversity and its spatial distribution^4^.

Introgressive hybridization is one of the most widespread forms of reticulate evolution and commonly occurs when recently diverged lineages regain geographical contact. Hybridization followed by repeated backcrossing transfers genetic material from one lineage into the genomic background of another. Because recombination fragments introgressed ancestry and selection influences its retention, different genomic regions may support conflicting genealogical relationships^5^. Introgression of mitochondrial genomes can additionally produce pronounced mito-nuclear discordance through mitochondrial capture^6^. Similar phylogenetic conflict can arise from incomplete lineage sorting (ILS), in which ancestral polymorphisms persist across successive divergence events and are stochastically sorted among descendant lineages^7^. Ancient introgression and ILS may consequently generate superficially similar distributions of discordant gene trees and can be difficult to distinguish using a small number of loci. At the geographical level, incomplete, uneven or spatially coarse sampling may further bias evolutionary inference by omitting admixed populations, peripheral remnants or the closest sampled relatives of extinct or unsampled donor lineages. Comprehensive geographical sampling is therefore essential for reconstructing diversification patterns in groups shaped by deep ancestral population structure^8^.

Butterflies provide some of the best-documented examples of introgression and ILS. Genome-wide analyses of *Heliconius* have revealed extensive exchange of ancestry across a rapid radiation, including the transfer of loci underlying wing-pattern mimicry and other adaptive traits^9,10^. More recently, multilocus introgression of traits related to colour pattern, wing shape, host-plant preference, pheromones and mate choice was shown to have played a central role in the origin and persistence of the hybrid species *H. elevatus*^11^. Both introgression and ILS are also evident in swallowtail butterflies. Whole-genome analyses of the Mediterranean *Papilio machaon* complex revealed clearer nuclear differentiation than was apparent from mitochondrial data, whereas genomic analysis of the hybrid zone between *P. syfanius* and *P. maackii* identified recent introgression and heterogeneous genetic exchange across the genome^12,13^. Together, these studies indicate that introgression and ILS are recurrent components of butterfly diversification, although their relative contributions may vary among phylogenetic depths, genomic regions and geographical scales.

The *Papilio bianor* species complex provides a particularly challenging system in which to evaluate these processes. Under the conservative taxonomic framework adopted here, the complex comprises three species with extensive but contrasting geographical distributions. *Papilio bianor* occurs broadly from the Himalaya and Indochina through eastern continental Asia and the peripheral islands of the southern Ryukyus; *P. dehaanii* is distributed mainly across the Japanese archipelago and adjacent parts of northeastern continental Asia; and *P. okinawensis* is largely restricted to the Amami and Okinawa island groups. Previous studies based on mitochondrial sequences or a limited number of mitochondrial and nuclear loci recovered deeply divergent lineages within the complex^14,15^. Mitochondrial ND5 analyses revealed similar geographical heterogeneity but provided limited resolution of population relationships^16^. Moreover, samples assigned to deeply divergent mitochondrial lineages were geographically intermingled, suggesting that their present distributions cannot be explained solely by simple, long-term geographical isolation. Instead, the observed pattern may record a temporally layered history involving ancient population structure, lineage turnover, mitochondrial capture, introgression and ILS.

This evolutionary complexity has contributed to persistent taxonomic instability. Western Himalayan *polyctor* and eastern *bianor* have alternatively been treated as separate species or as geographical forms within a broadly defined *P. bianor*, whereas *dehaanii* and *okinawensis* were historically classified as subspecies of *P. bianor* sensu lato. Molecular and taxonomic studies increasingly support the recognition of *P. dehaanii* and *P. okinawensis* as independent species, but the status of *polyctor*, Yaeyama *junia* and several other geographical forms remains unresolved^14,15,17^. Earlier multilocus analyses placed *polyctor* with *junia* and a Chinese sample in a deeply divergent lineage rather than recovering *polyctor* as sister to all other continental *P. bianor*^15^. Because most previous studies were based on mitochondrial sequences, a small number of nuclear loci or geographically restricted samples, their results could not adequately distinguish species divergence from ILS, introgression, mitochondrial capture or fragmented ancestry inherited from formerly widespread or unsampled populations. We therefore adopt a conservative three-species framework that recognizes *P. dehaanii* and *P. okinawensis* as independent species while provisionally retaining *polyctor* and *junia* as infraspecific taxa of *P. bianor*.

Advances in high-throughput sequencing now make it possible to evaluate these alternative evolutionary processes across entire genomes and densely sampled populations. Local phylogenies inferred from genomic windows can reveal variation in genealogical history among genomic regions, whereas summary-coalescent approaches can integrate these local trees while accounting for ILS^7,18^. Genome-wide SNP analyses, including principal component and ancestry analyses, D- and *f*-statistics, phylogenetic networks and explicit demographic models, can distinguish population structure from excess allele sharing and test alternative histories of divergence and admixture^19^. Combined with genomic scans of differentiation, nucleotide diversity and selection, these approaches can further determine whether introgressed ancestry is diffusely distributed or preferentially retained in regions associated with reproductive isolation and local adaptation^5^. Nevertheless, genomic resolution must be paired with geographically comprehensive sampling, because additional loci alone cannot recover evolutionary lineages or ancestral components that are absent from the sampled populations^8^.

Here, we analyse high-coverage whole-genome SNVs from 325 individuals representing geographically extensive sampling of *P. bianor*, *P. dehaanii* and *P. okinawensis*. We integrate concatenated and window-based phylogenomics, multispecies-coalescent inference, population-structure analyses, hierarchical tests of ILS and introgression, demographic modelling, and genome-wide analyses of differentiation and selection. Specifically, we quantify the relative contributions of ILS and gene flow across species, evolutionarily significant units and geographical populations; investigate the origin of mito-nuclear discordance; reconstruct the demographic history of the principal *P. bianor* lineages; and test whether geographically restricted populations retain ancestry from formerly widespread or unsampled lineages. We further assess how palaeoclimate, biogeography, admixture, demographic replacement, peripheral isolation and local adaptation jointly generated present-day diversity. By revealing how deeply differentiated ancestral populations contributed to subsequent lineage diversification, this study offers insight into how ancestral variation and historical gene flow jointly shape the rapid emergence of lineage diversity during the early stages of speciation.

## Results

### Chromosome Synteny and Reference Genome Annotation

Chromosome synteny analysis confirmed that Chr01 is the Z chromosome of *P. bianor* and revealed substantial chromosomal rearrangements between *P. bianor* and *P. xuthus* (Supplementary Fig. 1a). The genome annotation pipeline generated a comprehensive annotation set comprising 13,828 gene models and 12,819 CDS models (Supplementary Table S14). The majority of gene models were assigned to protein-coding loci (86.54%), followed by non-coding RNAs (9.57%) and pseudogenes (3.88%). Functional annotation assessment indicated that 35.63% of genes possessed confidently assigned gene names or symbols, while an additional 40.37% were supported by informative functional predictions. Approximately one quarter of genes (24.00%) remained without detectable functional annotation. Similar proportions were observed at the CDS level, with 38.19% of CDS models associated with known genes, 37.04% with predicted functions, and 24.77% lacking functional assignments.

### Variant Discovery

Genome-wide sequencing depth and coverage for each individual (Supplementary Table S1) are provided in Supplementary Table 1. Individuals with sequencing depth below 5× were excluded. We additionally applied lineage-specific coverage thresholds of 90% for *P. bianor* (excluding *P. b. polyctor* because of limited sample size), 85% for *P. dehaanii* and 80% for *P. okinawensis*. After quality filtering, we obtained high-quality genome-wide variant datasets from 325 individuals, including 319 ingroup samples (*P. bianor*, n = 166; *P. dehaanii*, n = 112; and *P. okinawensis*, n = 41) and six outgroup samples (*P. paris*, n = 1; *P. hermosanus*, n = 1; *P. maackii*, n = 3; and *P. krishna*, n = 1). Mean genome coverage and sequencing depth were 96.19% (86.22–98.12%) and 29× (7–113×) for *P. bianor*, 90.55% (87.34–92.95%) and 18× (8–37×) for *P. dehaanii*, 87.23% (83.85–91.20%) and 26× (11–84×) for *P. okinawensis*, and 75.76% (74.98–77.22%) and 24× (20–28×) for the outgroups.

Stringent filtering retained 47,828,665 high-confidence SNPs in dataset (a), with their genomic distribution shown in Supplementary Fig. 1b. Dataset (b) contained 49,039,202 variant (Supplementary Fig. 1c) and 106,547,748 invariant sites, 155,586,950 sites in total. The final mitochondrial genome alignment was 15,333 bp long.

### Population structure of the *P. bianor* complex

To characterize hierarchical genetic structure across the *Papilio bianor* species complex, we conducted phylogenetic reconstruction, principal component analysis, and ancestry inference based on genome-wide single-nucleotide polymorphism (SNP) data. The maximum-likelihood phylogeny based on genome-wide common variants (MAF ≥ 0.05) recovered three strongly differentiated clades corresponding to *P. bianor*, *P. dehaanii*, and *P. okinawensis* (Fig. 1c). Within *P. bianor*, Indochinese and eastern Chinese populations were partially intermingled but together formed a monophyletic group, which was subsequently sister to the Himalayan lineage, whereas the island lineage was positioned outside all continental lineages. Geographical populations were delineated based on preliminary population structure analyses and a morphological classification framework primarily informed by wing patterns; their abbreviations are provided in Table 1.

**Fig. 1.**
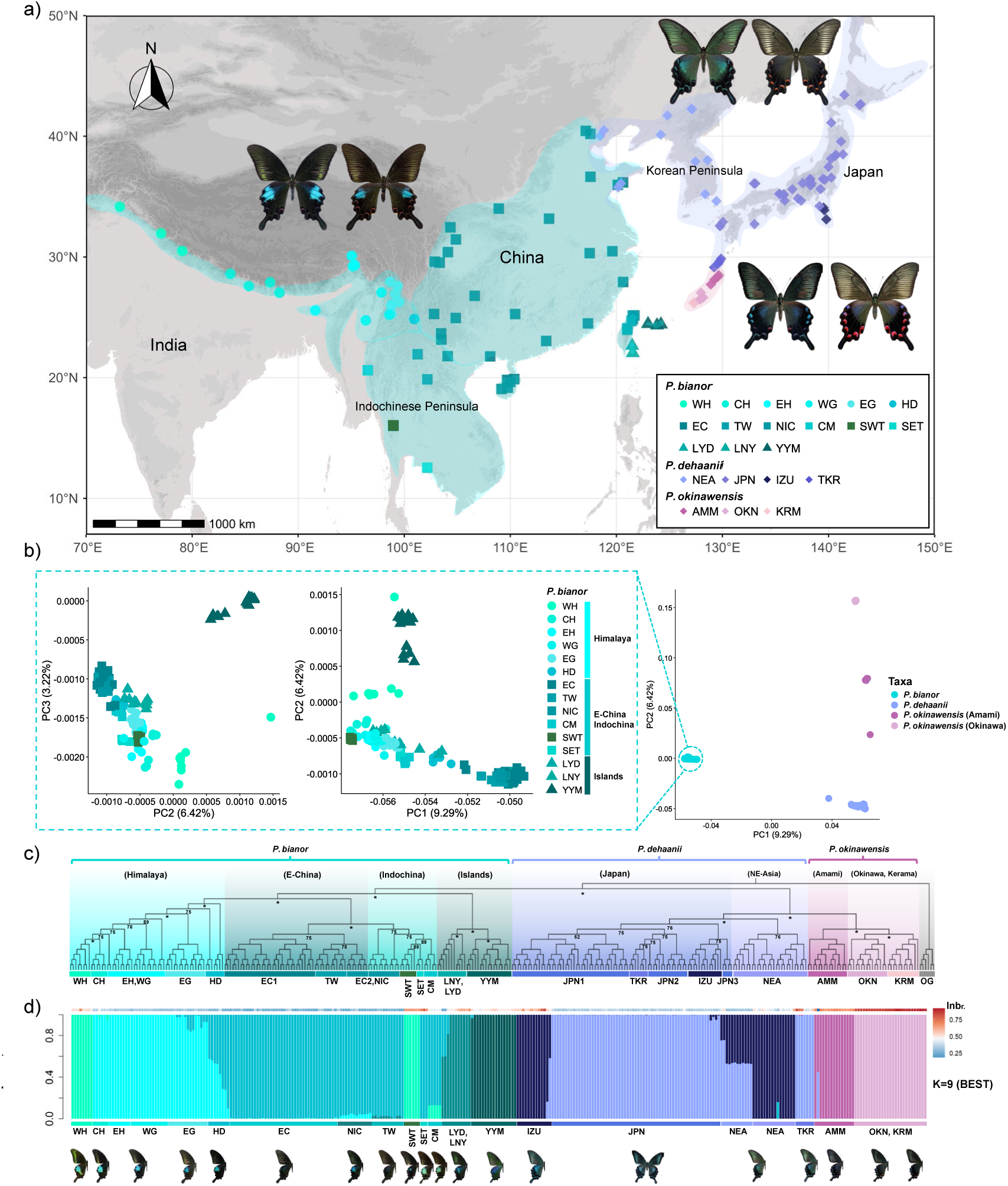
Sampling distribution and population structure of the *Papilio bianor* species complex and its closely related species, *P. dehaanii* and *P. okinawensis*. (a) Geographic distributions and sampling localities of *P. bianor* (*n* = 166), *P. dehaanii* (*n* = 112), and *P. okinawensis* (*n* = 41). (b) Principal component analysis of all ingroup samples based on dataset (a) retained after linkage-disequilibrium pruning at *r*² = 0.2, with an enlarged view showing the fine-scale genetic structure within *P. bianor*. (c) Maximum-likelihood phylogeny inferred in IQ-TREE from the complete dataset (a) with all 325 samples, including 319 ingroup and five outgroup individuals. Bootstrap support values from 1,000 replicates are shown for major nodes, together with the corresponding geographic groups, wing-pattern diversity, and individual inbreeding coefficients. (d) ADMIXTURE analysis of all ingroup samples based on the dataset (a) retained after linkage-disequilibrium pruning at *r*² = 0.2, showing the clustering solutions for *K* = 8, 9, and 10 and their correspondence with geographic population assignments. The lowest cross-validation error was obtained at *K* = 8.

**Table 1:**
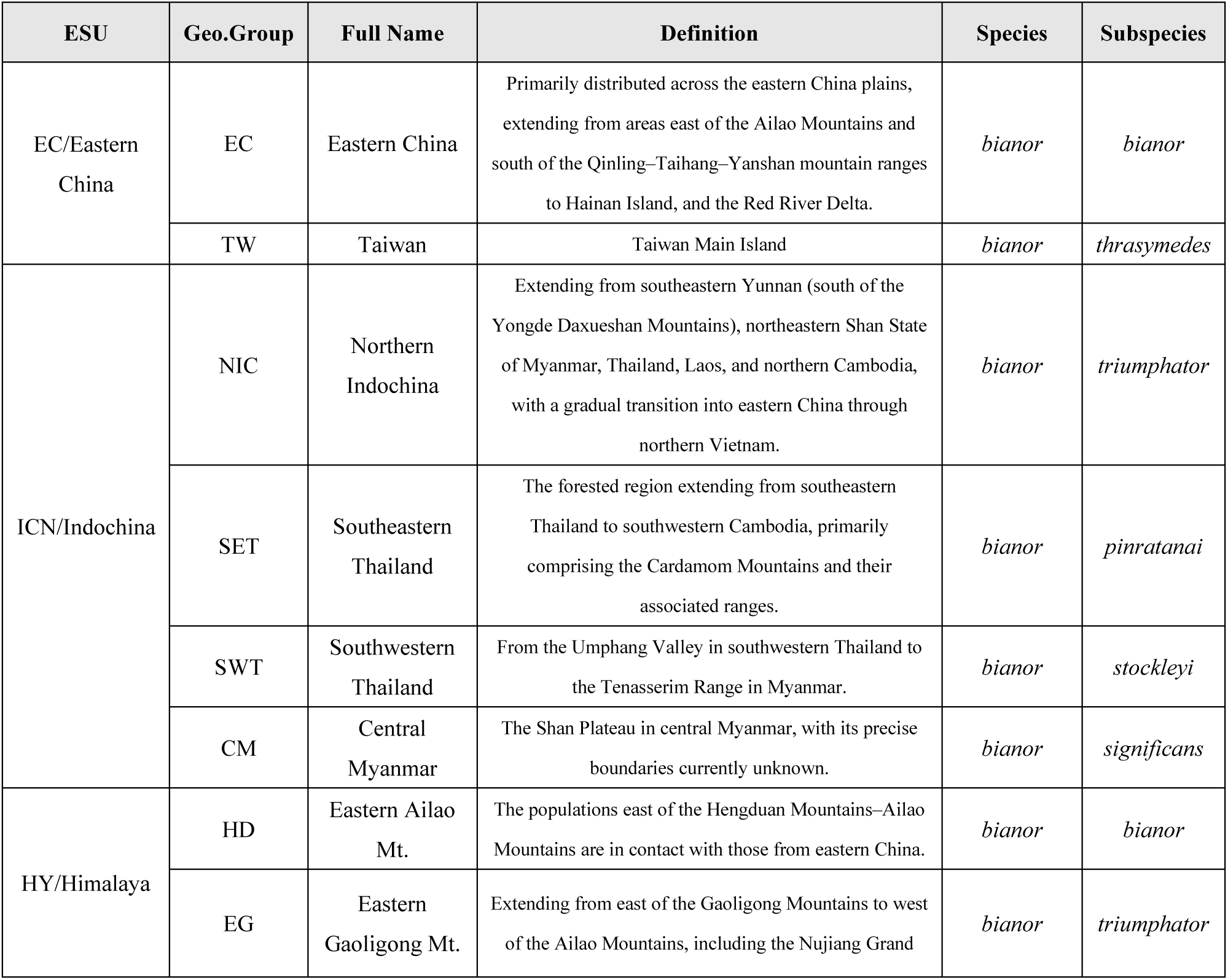
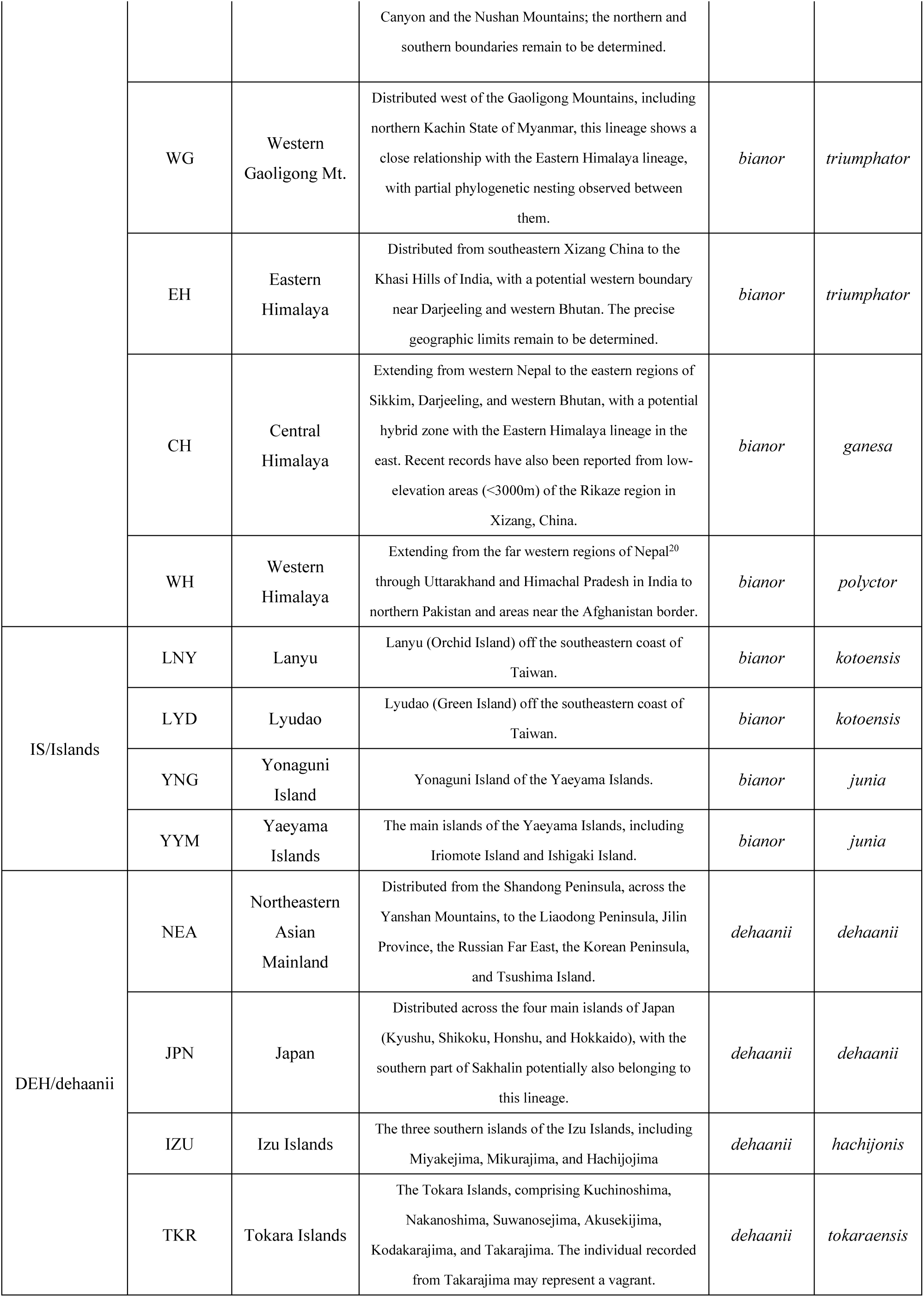
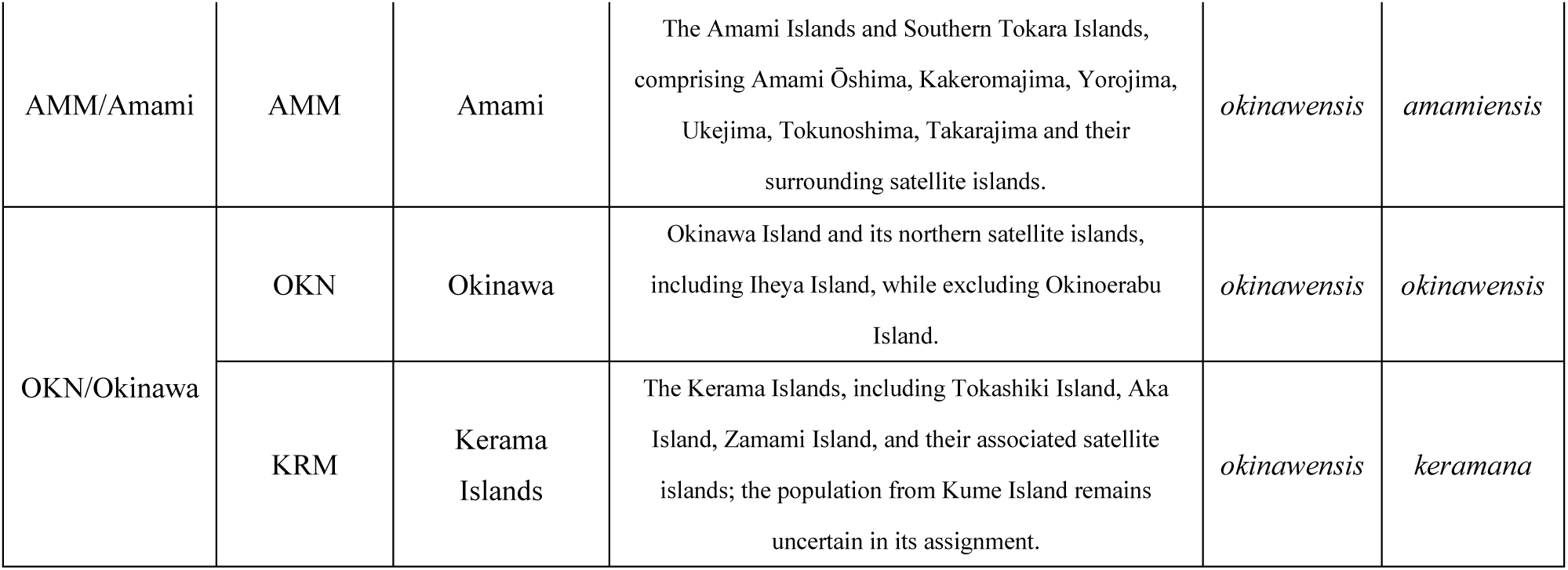
Geographic population delimitation, corresponding evolutionary significant units (ESUs), and currently recognized species and subspecies under the existing taxonomic framework.

At a finer geographical scale, LNY from Lanyu and LYD from Ludao formed a clade sister to YYM from the Yaeyama Islands, together constituting the island lineage. Within the Himalayan lineage, populations east of the Ailao Mountains diverged earliest, with major phylogeographical breaks occurring near the Gaoligong Mountains and between eastern Nepal and Darjeeling. CH from the central Himalaya and WH from the western Himalaya formed a supported clade, whereas populations from the eastern Himalaya exhibited a more complex and poorly resolved pattern involving multiple lineages. Eastern Chinese and Indochinese populations were not reciprocally monophyletic, with several regional populations showing intermingled phylogenetic relationships.

Principal component analysis based on all autosomal SNPs broadly matched the nuclear phylogeny. The first two principal components separated the three recognized species and explained 9.29% and 6.42% of the total variation, respectively (Fig. 1b). Within *P. okinawensis*, the Okinawa–Kerama and Amami groups formed distinct clusters, although one Amami individual occupied an intermediate position towards *P. dehaanii*. Most *P. dehaanii* individuals formed a compact cluster, except for one Qingdao individual slightly displaced towards *P. bianor*.

Within *P. bianor*, PCA revealed pronounced geographical genetic structure, with partial overlap among populations from adjacent regions. Along PC1, EC, TW, and NIC formed a relatively compact cluster, whereas Himalayan populations other than WH clustered together toward more negative PC1 values, with HD showing a gradual shift toward EC. SWT was clearly separated from the other Indochinese populations and occupied a position closer to the central Himalayan population CH in the PCA space. The island populations showed a distinct pattern: YYM was clearly separated from the continental populations along PC2, whereas LNY and LYD were positioned closer to the Himalayan populations and did not cluster with YYM. In contrast, WH did not cluster with the other Himalayan populations but was instead displaced toward YYM. PC3, which explained 3.22% of the total genetic variation, further resolved the genetic structure among geographical populations. Most notably, YYM and WH were both clearly separated from all other populations; however, unlike their relative proximity in the PC1–PC2 space, they were separated from each other along PC3 and occupied opposite ends of this axis. Most WH individuals were positioned closer to CH, whereas a single individual from Uttarakhand was distinctly displaced toward positive PC2 values (Fig. 1b).

ADMIXTURE further resolved these subdivisions, with K = 9 showing the lowest cross-validation error (CV error = 0.15986) and therefore being selected as the optimal number of ancestral components (Fig. 1d; Supplementary Fig. 4). At K = 9, the inferred ancestry components within *P. bianor* broadly corresponded to western Himalayan–southwestern Thailand, eastern Himalayan–Hengduan Mountains, eastern China–southeastern Thailand, and two island-associated components. WH and SWT exhibited nearly identical ancestry profiles and were dominated by the same major ancestry component. Among the Himalayan populations, EG contained a minor E-China-related ancestry component, whereas HD showed varying degrees of admixture between Himalayan- and E-China-related components; this mixed ancestry pattern was not observed in the more western Himalayan populations, including WG, EH and CH. EC, TW and NIC showed highly similar ancestry compositions, and SET displayed a closely related ancestry profile. Island populations were characterized by two distinct island-associated ancestry components, with island ancestry nearly fixed in YYM and predominant in LNY and LYD, although LNY and LYD also retained variable mainland-related ancestry. Within *P. dehaanii*, ADMIXTURE further resolved differentiation among regional populations, whereas the Amami and Okinawa–Kerama groups of *P. okinawensis* remained consistently distinct.

Pairwise Weir and Cockerham’s *F*ST revealed pronounced geographical heterogeneity and generally stronger differentiation on the Z chromosome than on the autosomes (Supplementary Fig. 2). Autosomal differentiation was low across the eastern Himalayan–Hengduan–eastern China–northern Indochina continuum, including EH–EG (FST = 0.016), EG–HD (0.013), EC–TW (0.016) and EC–NIC (0.007). Corresponding Z-chromosome values were generally higher but remained low among the most closely connected populations, including EH–EG (0.078), EG–HD (−0.004), EC–TW (0.024) and EC–NIC (0.008). WH was consistently differentiated from other Himalayan and eastern continental populations, with FST values of 0.145–0.221 on the autosomes and 0.217–0.416 on the Z chromosome. LNY and YYM were moderately differentiated from one another (autosomes, 0.288; Z chromosome, 0.400) and were genetically closer to EC, TW and NIC than to southern Indochinese populations. SET and SWT were strongly differentiated from most other *P. bianor* populations and from each other, with pairwise FST increasing from 0.543 on the autosomes to 0.668 on the Z chromosome, whereas CM showed intermediate differentiation.

Individual inbreeding coefficients showed a corresponding geographical pattern (Supplementary Table S3) (Fig. 1c). EC, TW, NIC and adjacent Himalayan–Hengduan populations generally had low-to-moderate values, whereas WH and island populations showed greater variation. The highest coefficients occurred in the geographically restricted Okinawa and Kerama populations of *P. okinawensis*, while most *P. dehaanii* populations had comparatively low values.

Overall, the nuclear phylogeny, PCA, ADMIXTURE, *F*ST and inbreeding coefficients revealed concordant genetic subdivision across the complex. Nuclear genomic analyses separated the three recognized species and identified geographical structure within them. Within *P. bianor*, analyses consistently distinguished a broadly connected continental core from strongly differentiated peripheral populations, particularly WH, the island lineage and southern Indochinese populations. Populations of *P. dehaanii* showed comparatively limited genomic differentiation and were treated as a single evolutionarily significant unit (ESU), DEH. By contrast, *P. okinawensis* comprised two distinct ESUs corresponding to Amami (AMM) and Okinawa–Kerama (OKN).

The complex was therefore summarized into seven ESUs: four within *P. bianor*—Himalaya (HY: WH, CH, EH, WG, EG and HD), eastern China (EC: EC and TW), Indochina (ICN: NIC, CM, SWT and SET) and Islands (IS: YYM, LNY and LYD)—one within *P. dehaanii* (DEH), and two within *P. okinawensis* (AMM and OKN) (Table 1). Additional geographical differentiation remained within WH, SWT, SET and individual island populations of *P. bianor*.

### Reticulate evolution across hierarchical levels

The 18,635 maximum-likelihood trees inferred from non-overlapping 20-kb autosomal windows revealed pervasive phylogenetic discordance across the *Papilio bianor* complex (Fig. 2a). ASTRAL recovered *P. bianor* as sister to the *P. dehaanii*–*P. okinawensis* clade and retained the island lineage outside the continental *P. bianor* lineages, although alternative quartet topologies occurred at substantial frequencies across many internal nodes. Within *P. bianor*, TWISST further revealed extensive variation among local genealogies (Fig. 2b). The most frequent topology (T1) grouped Indochina with the Himalaya, placed E-China as their sister lineage, and retained LNY and YYM as an island pair. T1 accounted for 11.93% of autosomal windows and increased to 14.54% on the Z chromosome. The alternative topology grouping Indochina with E-China (T2) was the second most frequent, accounting for 8.28% and 9.38% of autosomal and Z-linked windows, respectively. The third topology, grouping the Himalaya with E-China, occurred at similar frequencies on the autosomes (7.59%) and Z chromosome (7.62%). Two additional common topologies involving alternative placements of E-China and the island lineages each contributed 6.33–8.21% of Z-linked and 7.19–7.53% of autosomal windows. Thus, although no single genealogy dominated the genome, T1 was consistently the most frequent topology and showed a greater relative predominance on the Z chromosome.

**Fig. 2.**
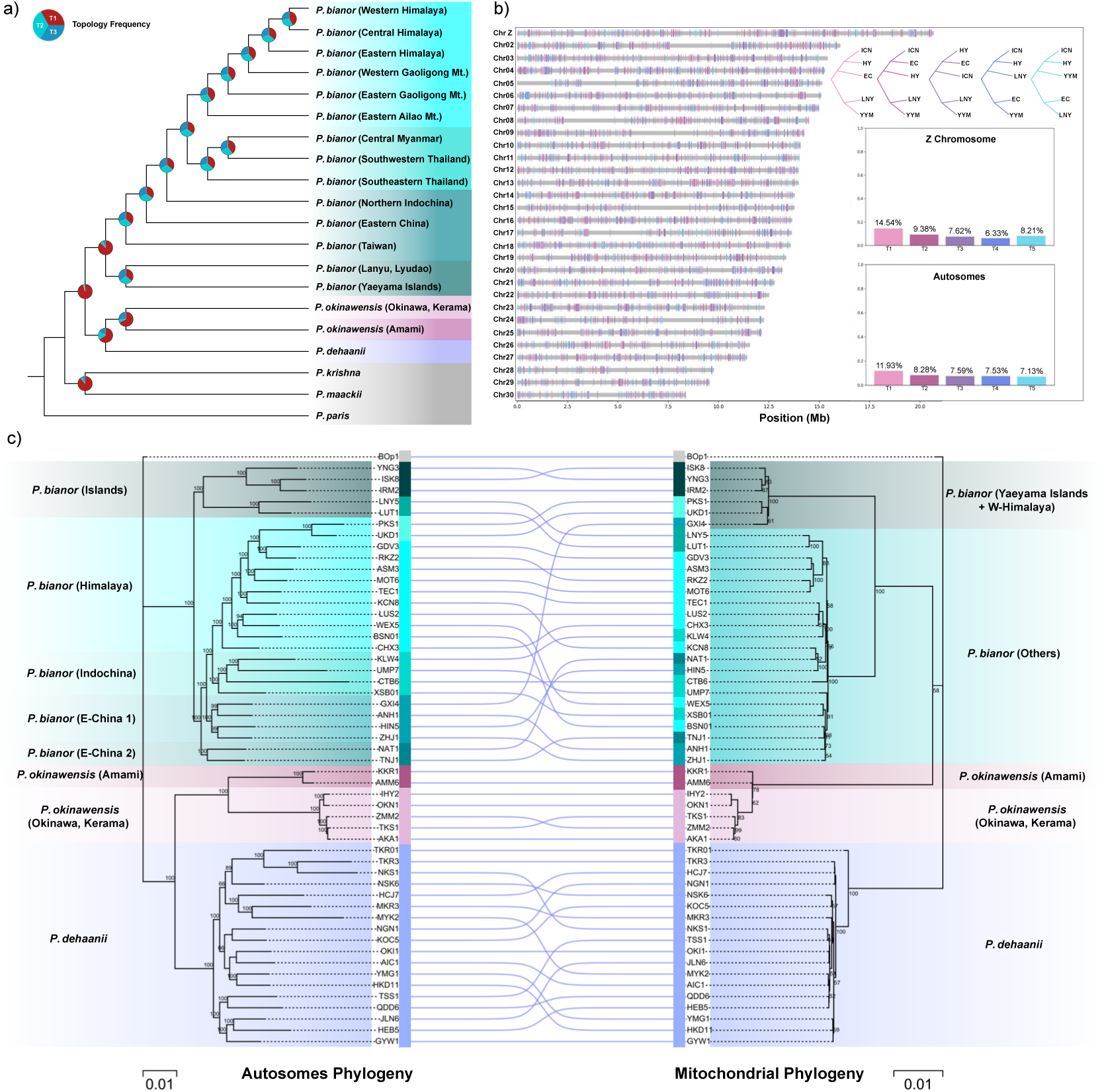
Genome-wide phylogenetic discordance and cytonuclear conflict in the *Papilio bianor* species complex and its closely related species, *P. dehaanii* and *P. okinawensis*. (a) ASTRAL species tree inferred from 18,635 maximum-likelihood trees generated from non-overlapping 20-kb autosomal windows. Pie charts at internal nodes show the relative frequencies of the three alternative quartet topologies (T1–T3), summarized using Astral-Hybrid, illustrating the extent of local gene-tree discordance across the species tree. (b) Genome-wide distribution of the five most frequent *P. bianor* topologies (T1–T5) identified by TWISST across the Z chromosome and 29 autosomes. Colored segments indicate 20-kb windows assigned to each of the five topologies, with the corresponding topology diagrams shown on the right. Bar plots summarize their proportions separately for the Z chromosome and autosomes. T1, which groups Indochina and the Himalaya as sister lineages, was the most frequent topology on both the Z chromosome (14.54%) and autosomes (11.93%), followed by T2 (9.38% and 8.28%, respectively); the remaining three major topologies occurred at lower frequencies. (c) Cytonuclear phylogenetic discordance among 53 representative samples. Maximum-likelihood trees were independently inferred from the autosomal SNP dataset and the 15,333-bp mitochondrial-genome alignment. Bootstrap support values based on 1,000 replicates are shown at internal nodes, and lines connect corresponding individuals between the two phylogenies. Shaded backgrounds denote major phylogeographic lineages. The most pronounced discordance involved the western Himalayan lineage (WH), which clustered with the other Himalayan populations in the autosomal phylogeny but with the Yaeyama Islands lineage in the mitochondrial phylogeny.

This nuclear topological heterogeneity was accompanied by pronounced mitonuclear discordance (Fig. 2c). The autosomal phylogeny placed WH within the Himalayan lineage and the Yaeyama populations within the island lineage, whereas the mitochondrial phylogeny grouped WH with the Yaeyama Islands. The remaining *P. bianor* populations formed a separate mitochondrial assemblage. Discordant placements were also observed within *P. dehaanii* and *P. okinawensis*, although the WH–Yaeyama association represented the most conspicuous conflict between the nuclear and mitochondrial phylogenies.

We next evaluated the relative contributions of incomplete lineage sorting (ILS) and introgression to genome-wide phylogenetic discordance using complementary approaches (Fig. 3). At the ESU level, Dsuite detected widespread allele-sharing signals within *P. bianor* and between *P. bianor* and its closely related species (Fig. 3a). Within *P. bianor*, the strongest signal involved the E-China and Islands lineages, whereas interspecific signals were most pronounced between *P. dehaanii* and the Amami lineage of *P. okinawensis*. f-branch analysis further localized these signals to specific evolutionary branches, identifying strong allele-sharing between E-China and Islands and between Amami and *P. dehaanii* (Fig. 3b,d).

**Fig. 3.**
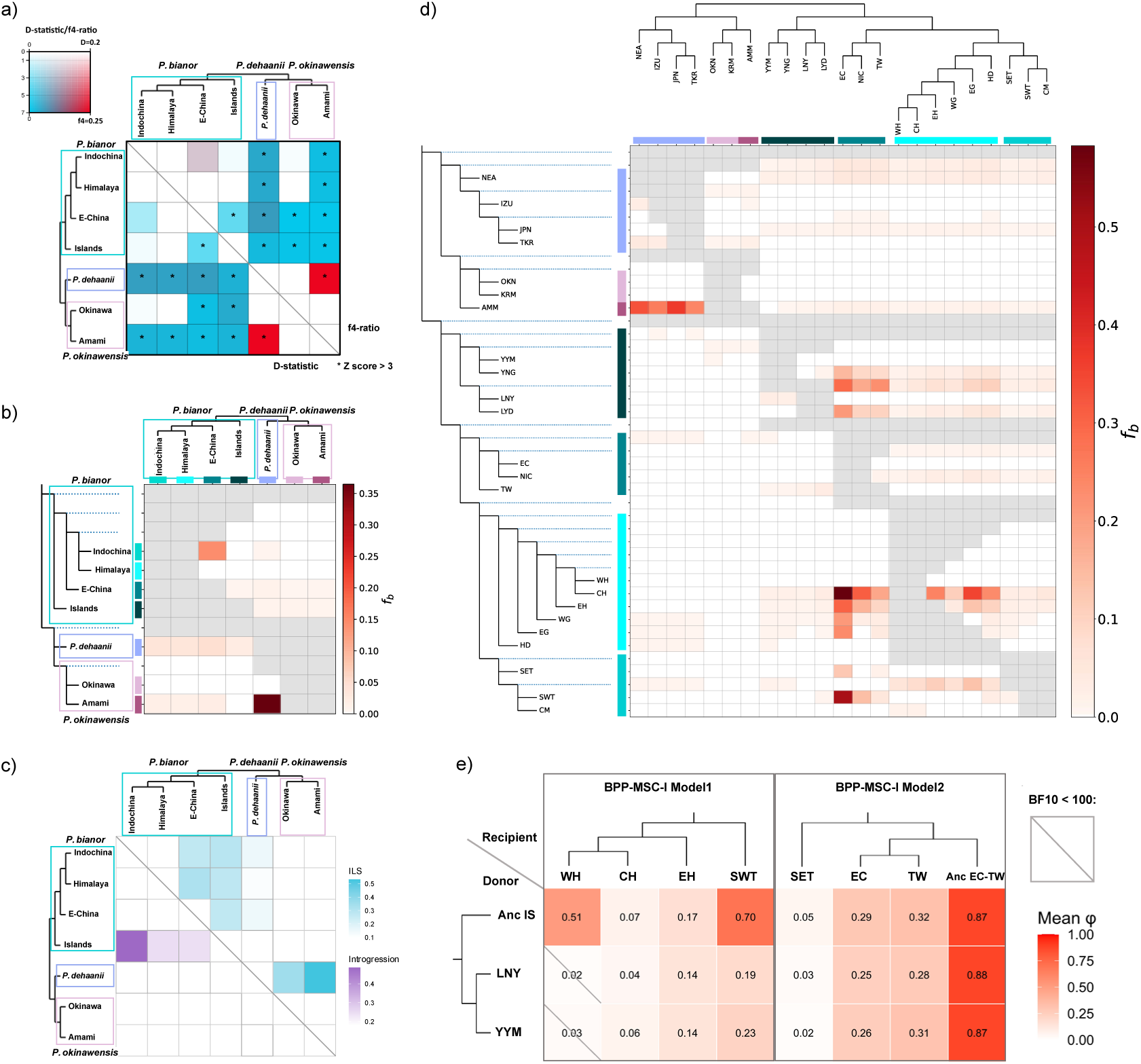
Genome-wide signatures of introgression, incomplete lineage sorting and ancestral ghost admixture in the *Papilio bianor* complex. (a) Patterson’s D-statistics and *f*4-ratios among the seven ESUs. Color intensity represents the magnitude of allele-sharing signals, and asterisks indicate significant signals (*Z* > 3). (b) f-branch analysis based on the species-tree topology showing the distribution and relative strength of gene-flow signals among ESUs. (c) Relative contributions of introgression and incomplete lineage sorting (ILS) estimated using QuIBL from genome-wide 20-kb window trees. (d) Population-level f-branch analysis showing heterogeneous allele-sharing patterns among geographical populations. (e) BPP MSC-I analyses testing ancestral island-related introgression scenarios. Heatmaps show mean introgression probabilities (φ) between donor lineages and recipient populations; only strongly supported models (BF₁₀ > 100) are shown. (f) HyDe estimates of parental contributions to the Eastern China lineage under alternative parental combinations involving mainland and island-related lineages. (g) GhostParser results identifying candidate sampled or unsampled donor lineages contributing to observed allele-sharing patterns. (h) TreeMix inference of migration events under the optimal model (*m* = 2), with migration edges indicating inferred historical gene flow among lineages.

QuIBL analyses distinguished discordance caused by ILS from introgression-associated signals (Fig. 3c). Among the four *P. bianor* ESUs, the strongest introgression component involved the Islands lineage and Indochina, with additional introgression signals involving Islands and both E-China and Himalaya. In contrast, discordance among continental lineages was predominantly attributed to ILS. Strong ILS signals were also detected among closely related lineages of *P. dehaanii* and *P. okinawensis*.

HyDe supported an admixed origin of the E-China lineage (Supplementary Fig. 6). Depending on the parental combination tested, E-China received a predominant continental ancestry component together with a substantial island-related contribution. When Indochina and Islands were specified as parental lineages, the estimated contributions were 63.6% and 36.4%, respectively. When Himalaya and Islands were used, the corresponding contributions were 71.78% and 28.22% (Supplementary Tables S10b, S10c). These results consistently supported the presence of island-related ancestry in E-China.

BPP MSC-I analyses further identified multiple strongly supported island-related introgression scenarios (Supplementary Table S12b) (Fig. 3e). The strongest signal involved introgression from an ancestral island-related lineage into the common ancestor of the Eastern China–Taiwan lineage (Anc EC-TW), with a mean φ value of 0.87 (BF₁₀ > 100). Similar high contributions were also inferred from extant island lineages, including LNY (φ = 0.88) and YYM (φ = 0.87), indicating substantial island-related ancestry in the ancestor of the Eastern China–Taiwan lineage. Additional strongly supported introgression from the ancestral island lineage was detected into WH (φ = 0.51) and SWT (φ = 0.70), whereas SET showed consistently weak contributions from all island-related donors (φ = 0.02–0.05). Across most Himalayan and Indochinese populations, contributions from extant island lineages were generally lower than those from the ancestral island lineage. These results indicate that island-related ancestry was incorporated into continental populations through multiple temporally distinct introgression events, with ancestral island-related lineages representing the major source of island ancestry in several peripheral continental populations.

TreeMix identified two migration edges (m = 2), including a weak migration event from an ancestral *P. okinawensis*-related lineage into WH and a stronger migration event from the Amami lineage into *P. dehaanii* (Supplementary Fig. 9). However, BPP MSC-I analyses did not support elevated interspecific introgression specifically into WH, suggesting that the WH-related signal may represent ancient or unsampled ancestry rather than recent gene flow.

Together, these results indicate that phylogenetic discordance in the *P. bianor* complex reflects a combination of ancient incomplete lineage sorting and recurrent introgression involving ancestral island-related lineages.

### Demographic history of the *P. bianor* complex

To reconstruct demographic history across the four *P. bianor* ESUs, we analysed representative geographical populations using PSMC, fastsimcoal2 and BPP. PSMC trajectories spanning approximately 10 ka to 5 Ma revealed shared demographic fluctuations among populations, with lineage-specific differences in the magnitude and timing of population changes (Fig. 4). Most *P. bianor* populations reached their maximum effective population sizes (*N*e) before the Penultimate Glaciation (130–300 ka), followed by declines of varying intensity. Some populations subsequently entered relatively stable plateau-like phases, whereas others continued to decline without clear recovery. These demographic transitions broadly corresponded to major Middle and Late Pleistocene climatic fluctuations.

**Fig. 4.**
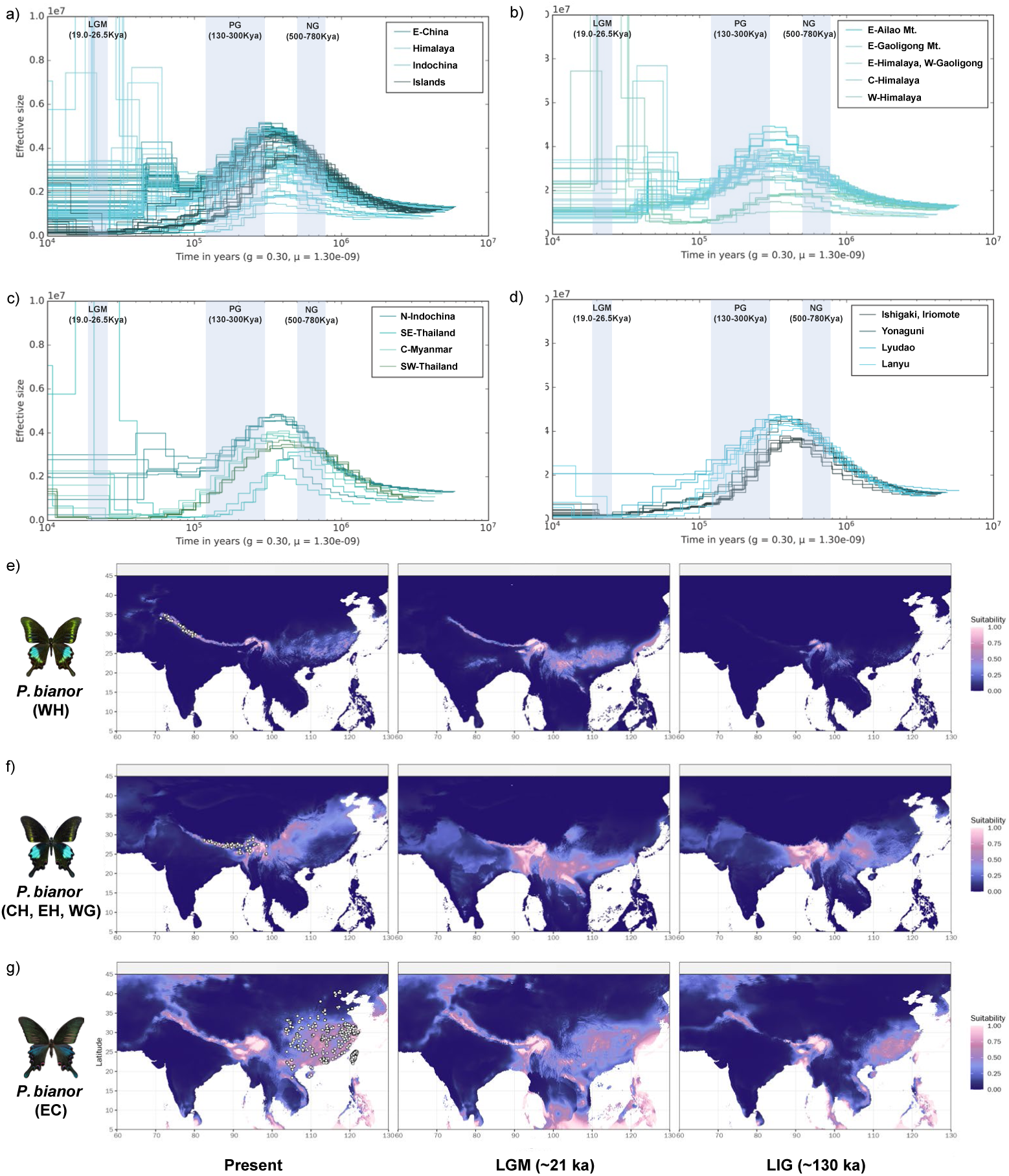
Demographic histories and historical distribution dynamics of *Papilio bianor* inferred using PSMC and species distribution models. Individual-based pairwise sequentially Markovian coalescent (PSMC) analyses were conducted for *P. bianor* (*n* = 100). The inferred effective population-size trajectories are shown together with three major glacial intervals: the Last Glacial Maximum (LGM), the Penultimate Glaciation (PG), and the Naynayxungla Glaciation (NG). (a) Demographic histories of the four evolutionarily significant units (ESUs) within *P. bianor*. (b) Demographic histories of geographic populations within the *P. bianor*–Himalaya lineage. (c) Demographic histories of geographic populations within the *P. bianor*–Indochina lineage. (d) Demographic histories of geographic populations within the *P. bianor*–Islands lineage. (e–g) Species distribution models showing temporal changes in potential distributions of three major continental lineages of *P. bianor*: western Himalaya (WH), eastern Himalaya–Hengduan Mountains (CH, EH and WG), and Eastern China (EC). Predicted habitat suitability was projected onto present-day climate conditions, the Last Glacial Maximum (∼21 ka), and the Last Interglacial (∼130 ka). Suitability values are represented by a continuous colour scale from low (dark blue) to high (pink). White circles indicate occurrence records used for model construction. Each row represents one lineage, and columns correspond to different climatic scenarios.

**Fig. 5.**
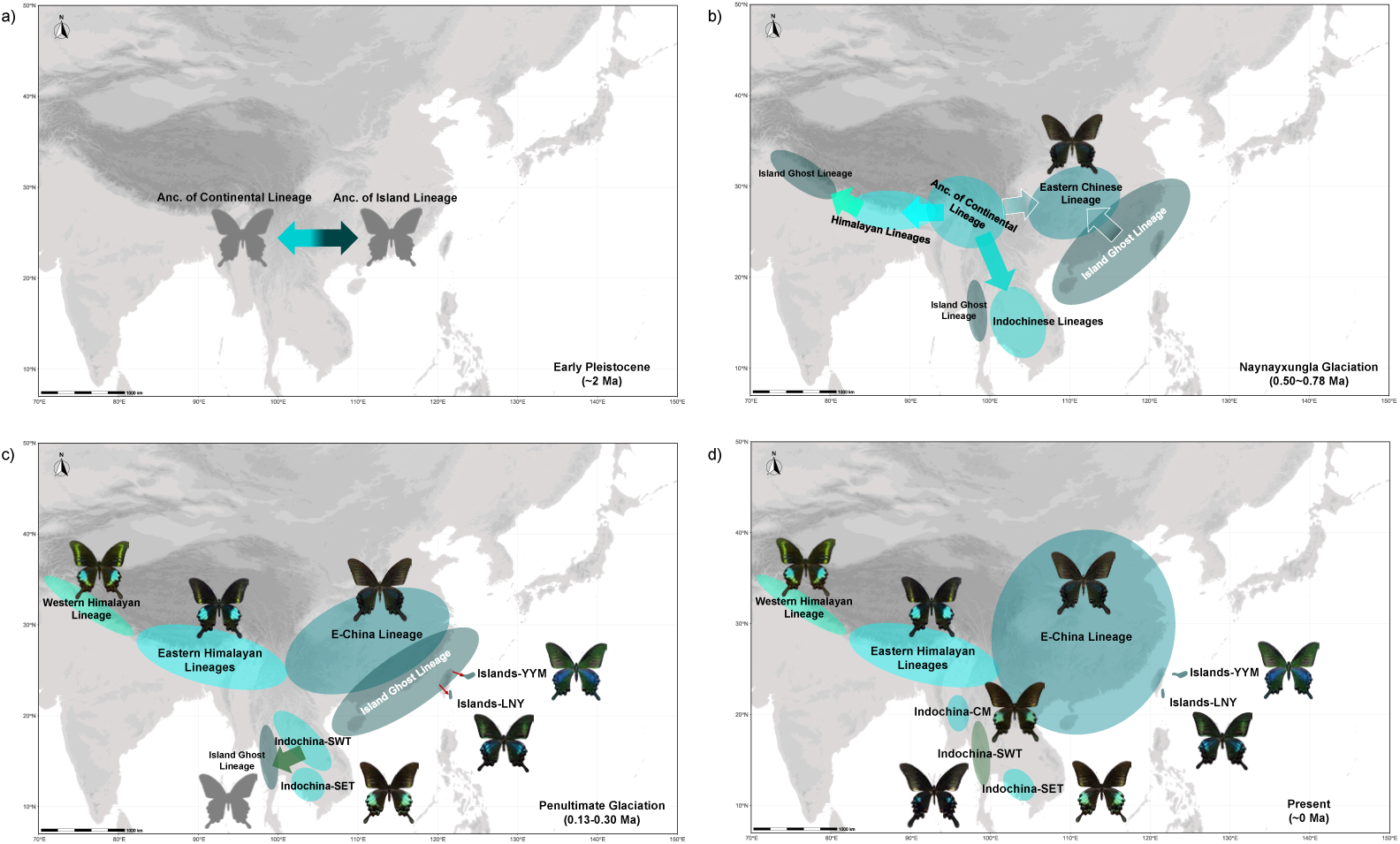
Diversification history and evolutionary dynamics of *Papilio bianor*. The best-supported demographic scenario inferred using fastsimcoal2 illustrates the formation and subsequent evolutionary dynamics of the four major evolutionarily significant units (ESUs) within *P. bianor*. The model summarizes mean divergence times, effective population sizes (*Ne*), migration rates among major lineages, and ancestry contributions associated with the origin of the Eastern China lineage. The evolutionary history of *P. bianor* involved an early divergence between island-related and continental ancestral lineages at approximately 1.86 Ma. The continental lineage subsequently underwent diversification during the Middle Pleistocene. The Eastern China lineage originated at approximately 0.66 Ma through admixture between the ancestral Himalaya–Indochina lineage and an island-related lineage, receiving approximately 37.92% and 62.08% ancestry contributions, respectively. The Himalaya and Indochina lineages subsequently diverged at approximately 0.119 Ma, indicating that their differentiation postdated the formation of the Eastern China lineage. Following its formation, the Eastern China lineage maintained asymmetric gene flow with neighbouring lineages, particularly stronger gene flow from the ancestral mainland lineage into Eastern China and from Eastern China into the Islands lineage. The Islands lineage experienced a pronounced demographic reduction and retained the smallest effective population size among the four major lineages. Together with BPP MSC-I analyses identifying multiple strongly supported ancestral island-related ghost introgression scenarios, these results indicate that diversification of *P. bianor* was shaped by sequential lineage divergence, secondary admixture involving ancestral island-related sources, and persistent asymmetric gene exchange.

Within *P. bianor*, demographic trajectories differed substantially among regional lineages. Eastern China and eastern Himalayan populations experienced declines after their demographic peaks but retained relatively large *N*e values or developed plateau-like trajectories (Fig. 4a,b). In contrast, the Islands lineage and the southwestern and southeastern Thailand populations continued to decline after their respective peaks, without evident demographic recovery (Fig. 4a,c). Within the Yaeyama Islands, Iriomote and Ishigaki maintained consistently larger *N*e values than the more geographically isolated Yonaguni population (Fig. 4d). More broadly, populations exhibiting stronger allele sharing with other lineages in f-branch analyses generally showed larger effective population sizes than their more isolated sister populations.

To resolve the formation of the four major *P. bianor* lineages, we next compared bifurcating with continuous gene flow and hybrid-origin demographic scenarios using fastsimcoal2 (Supplementary Fig. 13). The best-supported model inferred that the Islands lineage first diverged from the ancestral continental lineage at approximately 1.86 Ma (95% confidence interval, 1.82–1.90 Ma). The E-China lineage subsequently originated through admixture at approximately 0.66 Ma (0.61–0.72 Ma), involving the ancestral Himalaya–Indochina lineage and the Islands lineage, with estimated ancestry contributions of 37.92% and 62.08%, respectively (Fig. 4). The Himalaya and Indochina lineages diverged later, at approximately 0.119 Ma (0.112–0.123 Ma) (Supplementary Fig. 14; Supplementary Table S13).

Following the formation of E-China, gene flow with neighboring lineages was strongly asymmetric (Fig. 4). Migration from the Islands into E-China was highest (7.08 × 10⁻⁶), exceeding the reverse rate by more than an order of magnitude (1.39 × 10⁻⁷). Gene flow from Indochina into E-China was also stronger than in the opposite direction (2.22 × 10⁻⁶ versus 4.25 × 10⁻⁷), whereas exchange between the Himalaya and E-China was weaker and biased toward E-China (1.87 × 10⁻⁶ from the Himalaya into E-China versus 5.16 × 10⁻⁷ in the reverse direction). Contemporary effective population sizes differed markedly among lineages, with E-China showing the largest estimate (*N*e = 5,937,894; 95% confidence interval, 5,936,676–5,938,247), followed by the Himalaya (2,852,389; 2,707,296–2,988,287), Indochina (635,952; 620,848–651,789), and the Islands (184,663; 180,058–188,205). The Islands lineage declined from an ancestral *N*e of 311,523 (127,144–536,290), whereas the ancestral continental population increased from an estimated *N*e of 64,127 (10,651–155,624) before the initial divergence to 582,572 (248,014–874,978) and subsequently 424,824 (343,486–502,068) before the Himalaya–Indochina split.

Independent BPP MSC-I analyses provided a temporal framework for the three strongly supported ancestral island-related introgression events (Supplementary Fig. 15). Introgression from an unsampled ancestral island lineage into the common ancestor of EC and TW showed the highest estimated strength (mean φ = 0.87, BF₁₀ ≫ 100). In this model, the ghost donor lineage diverged from the lineage leading to extant LNY and YYM at approximately 0.83 Ma (95% HPD, 0.63–1.05 Ma), followed by introgression into the EC–TW ancestral lineage at approximately 0.78 Ma (0.52–1.04 Ma). A second strongly supported event involved introgression from an ancestral island ghost lineage into WH (φ = 0.51, BF₁₀ ≫ 100), for which the ghost lineage diverged at approximately 1.90 Ma (1.45–2.39 Ma) and introgression was dated to approximately 0.56 Ma (0.40–0.73 Ma). The third event involved introgression into SWT (φ = 0.70, BF₁₀ ≫ 100), with the ghost donor lineage diverging at approximately 1.27 Ma (0.96–1.61 Ma) and the introgression event occurring substantially later, at approximately 0.16 Ma (0.11–0.21 Ma). Together, these MSC-I models recovered substantial ancestral island-related introgression into geographically distinct continental lineages, with estimated introgression times ranging from approximately 0.78 to 0.16 Ma.

Together, these analyses indicate that *P. bianor* underwent early differentiation between island-related and continental lineages, followed by later diversification of the Himalayan and Indochinese lineages. The E-China lineage was formed by admixture between the ancestral Himalaya–Indochina lineage and island-related ancestry at approximately 0.66 Ma and subsequently experienced week gene flow with neighboring lineages. BPP MSC-I analyses further identified ancestral island-related introgression into the EC–TW ancestor, WH and SWT at different times during the Pleistocene. These demographic and reticulation events collectively shaped the present geographical structure of genomic diversity within the complex.

### Palaeodistribution Modelling

Palaeodistribution modelling revealed contrasting historical range dynamics among major *P. bianor* lineages (Fig. 4e, f, g). The western Himalayan lineage (WH) exhibited a restricted distribution under current climatic conditions, with suitable habitats mainly confined to the western Himalaya. During the Last Glacial Maximum (LGM, ∼21 ka), suitable areas expanded eastward across the Himalayan region, whereas during the Last Interglacial (LIG, ∼130 ka), suitable habitats contracted and were largely restricted to western Himalayan regions.

The eastern Himalayan lineage (CH, EH and WG) showed a broader distribution pattern. Current suitable habitats were concentrated in the eastern Himalaya and Hengduan Mountains, while LGM projections indicated a substantial expansion toward southwestern China and northern Indochina. During the LIG, suitable areas remained extensive but were reduced relative to the LGM, with persistence mainly in the eastern Himalayan–Hengduan region.

The Eastern China lineage (EC) maintained the broadest potential distribution among the three lineages. Under present climatic conditions, suitable habitats extended across eastern China, southern China, Taiwan and adjacent regions. During the LGM, suitable areas expanded southward and formed a continuous distribution across southern China and northern Indochina. The LIG projection showed a moderately reduced but still extensive suitable range concentrated in eastern and southern China. Overall, the SDM results indicate that all three continental lineages experienced glacial-period range expansion and interglacial contraction, but differed in the extent and geographic direction of historical range shifts. The WH lineage showed the strongest range restriction, whereas EC maintained the largest and most persistent potential distribution.

## Discussion

### Ancestral variation and ancient admixture shaped early lineage diversification

The *Papilio bianor* complex illustrates how ancestral genetic variation was retained, recombined, and transformed into regional diversity during the early stages of lineage divergence. Our results support a temporally layered evolutionary history in which early geographic differentiation established the major lineage structure, ancestral polymorphism and ancient admixture jointly shaped genomic composition, and subsequent ecological differentiation and regional isolation further reinforced lineage divergence. Previous studies based primarily on a limited number of molecular markers identified deeply divergent lineages within *P. bianor* but did not resolve the population structure of the continental lineages, precluding explicit links between ancestral structure and lineage diversification^14,15^. Our whole-genome analyses further reveal that admixture among ancestral lineages was an integral component of this diversification process.

The best-fitting fastsimcoal2 model dated the island–continental split to approximately 1.86 Ma and inferred an admixed origin of the Eastern China lineage at approximately 0.66 Ma, with 37.92% ancestry from the ancestral Himalaya–Indochina population and 62.08% from an island-related lineage. Whereas fastsimcoal2 dated the Himalaya–Indochina split to approximately 0.12 Ma, BPP MSC-I consistently recovered earlier, similar divergence times across introgression scenarios, supporting concentrated diversification of multiple lineages in both regions during the third-to-last glacial period. This difference may reflect the scale of population grouping: our fastsimcoal2 models treated each region as a single population, whereas BPP MSC-I retained pronounced population structure within regions and jointly modelled divergence, ILS, and introgression. The finer population resolution and consistent timing across scenarios strengthen support for this earlier phase of rapid diversification.

Multiple methods recovered both continental and island-related ancestry in the Eastern China lineage. HyDe estimated a greater continental contribution under configurations using extant parental lineages, whereas fastsimcoal2 and the initial BPP MSC-I analyses recovered a greater island-related contribution under ancestral admixture scenarios. These estimates draw on site patterns, site frequency spectra, and multilocus coalescent histories, respectively, and correspond to different parental configurations and historical parameters^21–23^. Their shared finding strongly supports an admixed origin of the Eastern China lineage, with ancient admixture contributing to its formation. Expanded-population BPP MSC-I analyses also recovered island-related introgression into the Eastern China ancestor (AncECTW), with a more recent inferred event and a lower island-related contribution (Supplementary Fig. 12a; Supplementary Table S12b). Together with the ranking of ancient and recent introgression as the best-fitting and second-best fastsimcoal2 models, respectively, these results motivate a hypothesis of temporally layered admixture involving more than one episode of island-related gene flow. As the most widely distributed and highly connected population in the complex, Eastern China provides a setting in which successive contacts could have involved different island-related donors, making its admixture history particularly complex. Explicit tests of multiple introgression episodes and alternative donor lineages offer a promising route to resolving this history. A related example occurs in *P. appalachiensis*, which combines genetic contributions from *P. glaucus* and *P. canadensis* with an ecological combination of mimicry and traits associated with cooler habitats^24^. In this study, the Eastern China lineage exhibited wing-surface patterning similar to that of the Yaeyama lineage, characterized by the absence of dense metallic-blue markings, while its wing shape resembled that of many continental lineages. This combination of traits was likely retained under long-term ecological pressures and natural selection, facilitating adaptation to the plains of eastern China.

Differential sorting of ancestral polymorphism accounts for extensive genetic sharing among continental lineages. Closely spaced divergence events allow ancestral alleles to persist across successive splits, producing genealogies that differ from the species tree^7^. Our topology analyses, PhyTop, and QuIBL identified ILS as the principal source of genealogical discordance among continental lineages, whereas stronger introgression signals involved continental and island-related lineages. Early diversification therefore involved both the redistribution of ancestral variation and the incorporation of ancestry from other lineages.

PSMC trajectories further reveal the temporal and spatial overlap of these processes. Regional curves are broadly similar in the deeper past but subsequently diverge in peak magnitude, the extent of decline, and later trajectories, with marked transitions also occurring within Himalaya and Indochina (Fig. 4a–d). Rapid diversification left descendant populations with extensively shared ancestral genealogies, while admixture between deeply divergent continental and island-related lineages brought genomic segments with different coalescent depths into the same genomes. Together with our divergence and introgression results, this process provides a coherent explanation for the combination of similarity, differentiation, and gradual transitions among regional PSMC trajectories.

### Peripheral populations retain distinct patterns of ancient ancestry

Western Himalaya (WH) and southwestern Thailand (SWT) belong to their respective continental nuclear lineages while showing distinctive genetic compositions in PCA and ADMIXTURE analyses. BPP MSC-I recovered significant island-related introgression signals involving the Eastern China–Taiwan ancestor, WH, and SWT; in WH and SWT, signals were particularly strong under ancestral donor scenarios relative to corresponding extant island donor scenarios. These signals persisted under expanded population sampling. For WH, the inferred timing and ancestry contribution remained broadly consistent across analyses, providing strong support for introgression from an island-related ghost lineage. SWT likewise showed strong evidence of island-related ancestry, with its estimated contribution varying substantially between analyses (Supplementary Fig. 12b, c; Supplementary Table S12b). This consistent detection identifies island-related ancestry as a component of SWT evolution, while its magnitude remains a target for further demographic modelling. These results support a contribution of island-related ancestry to evolution across multiple continental regions. Introgression involving deep ancestral branches in *Heliconius* likewise demonstrates the persistence of early genetic exchange in descendant genomes^23^.

The nuclear placement and mitochondrial affinity of WH reveal different components of this history. Its nuclear genome consistently places it within the Himalayan lineage, whereas its mitochondrial genome shows affinity with Yaeyama. Together with the ancestral introgression inferred by BPP, this pattern supports a role for ancient island-related ancestry in WH evolution. Similarly, admixed regional taxa in the *P. machaon* group retain particular mitochondrial lineages alongside distinct nuclear genetic compositions^25^. These findings illustrate how genomic compartments can differentially preserve the signatures of historical admixture.

Southeastern Thailand (SET) presents a further hypothesis for investigation. Its high FST relative to SWT highlights marked differentiation within Indochina and, together with the introgression signals recovered under alternative MSC-I configurations, motivates tests of a contribution from an external ghost lineage. Future comparisons of demographic models incorporating isolation, drift, and alternative ghost donors could clarify whether external introgression contributed to the distinctive ancestry of SET.

Ancient ancestry sharing and subsequent regional differentiation form successive parts of this process. Ancestral admixture established genetic connections among regions, after which isolation, population contraction, and drift altered ancestry frequencies and generated distinctive peripheral populations. The patterns in WH and SWT support this sequence, with present-day differentiation recording both early genetic connections and subsequent regional demographic histories.

### Ecological differentiation and Geographic isolation help maintain regional differentiation

Geographic isolation and ecological differences appear to maintain lineage diversity within the *P. bianor* complex. Admixture between Eastern China and Eastern Himalayan populations is concentrated around the Hengduan Mountains, whereas populations farther west retain distinctive ancestry. Species distribution models indicate relatively low habitat suitability for the Eastern China population in this region under both present and Last Glacial Maximum conditions, suggesting that habitat discontinuity has restricted connectivity over time. Prolonged isolation would also have strengthened genetic drift, particularly in WH, SWT, southeastern Thailand, and island populations, whose demographic histories show stronger contractions than those of eastern continental populations. Together, restricted connectivity and contrasting demographic histories help explain the persistence of regional genomic differences. A comparable pattern occurs in the closely related *P. syfanius* and *P. maackii*, which predominantly occupy highland and lower-elevation habitats, respectively, and remain genomically differentiated despite gene flow in their contact zone^12^.

Localized gene flow, divergent demographic histories, and relatively stable differences in wing pattern together suggest that speciation may be underway among some regional lineages. Their continued differentiation despite occasional genetic exchange is consistent with geographic isolation and ecological differences contributing to the maintenance of lineage boundaries. Although reproductive isolation has not been systematically assessed, these findings identify regional lineages at potentially different stages of divergence. The *P. bianor* complex thus provides evidence for a progression from ancestral admixture and rapid diversification to the maintenance of regional diversity through prolonged isolation, drift, and ecological differentiation.

## Methods

### Samples and whole-genome sequencing

We included 325 adult butterfly tissue samples or publicly available sequencing datasets from *Papilio bianor*, *P. dehaanii* and *P. okinawensis*, covering all recognized subspecies and several geographically defined populations without formal subspecific designation (Supplementary Table S1). Three additional species, *P. paris*, *P. maackii* and *P. krishna*, were included as outgroups (Supplementary Table 1).

Samples were collected between the late 1980s and 2025. Adult butterflies were collected in the field using hand nets or reared from eggs under laboratory conditions. Additional specimens were obtained from commercial suppliers and breeding facilities holding valid permits for collection, husbandry and importation, or were donated as dried material by private collectors. We also retrieved 12 publicly available whole-genome resequencing datasets from NCBI. For populations currently subject to restrictions on wild collection, only specimens collected before the relevant regulations came into force or individuals derived from captive-breeding programmes, including *P. dehaanii tokaraensis*, were included.

Genomic DNA was extracted from frozen tissue using a CTAB protocol. Samples were homogenized in liquid nitrogen, incubated in 2× CTAB buffer at 65 °C, extracted with phenol/chloroform/isoamyl alcohol, treated with RNase A and precipitated with isopropanol or ethanol. DNA pellets were washed with 80% ethanol and dissolved in nuclease-free water. Sequencing libraries were prepared by ultrasonic fragmentation, end repair and ligation of MGIEasy PF adapters, followed by magnetic-bead purification, PCR amplification and a second purification step. Fragments with an insert size of approximately 450 bp were selected by agarose-gel electrophoresis. Libraries were assessed using an Agilent Bioanalyzer and quantified by PicoGreen assay, and only libraries showing a single fragment peak, no detectable adapter dimers and concentrations of at least 2 nM were retained. Qualified libraries were pooled equimolarly, denatured to single-stranded DNA and sequenced on the DNBSEQ-T7 platform using 150-bp paired-end reads. Whole-genome resequencing was conducted at approximately 10–50× depth, primarily using leg tissue. Older dried specimens were generally sequenced at depths exceeding 30×. Genome-wide sequencing depth and coverage were calculated using PanDepth v2.26^26^.

### Reference genomes and variant calling

The chromosome-level nuclear reference genome of *P. bianor* (GCA_040363705.1)^27^ was downloaded from NCBI. Scaffolds not assigned to chromosomes were removed, leaving 30 nuclear chromosomes. Because this assembly lacks sex-chromosome annotations, we additionally downloaded the annotated genome of *P. xuthus* (GCA_036365505.1) and conducted chromosome-synteny analysis using NGenomeSyn v1.43^28^ to identify the Z chromosome (Supplementary Fig. 1). The mitochondrial reference genome of *P. bianor* (KF859738.1)^29^ was also retrieved from NCBI and used to reconstruct population-level mitochondrial sequences.

Transcriptomic reads and orthologous protein sequences used for genome annotation were downloaded from NCBI BioProject PRJNA1009405. Gene structures in the *Papilio bianor* reference genome (GCA accession number) were annotated using EGAPx v0.5.0 (NCBI; https://github.com/ncbi/egapx). Pseudogenes were identified with EasyPseudogene^30^, tRNAs with tRNAscan-SE v2.0.12^31^, and other non-coding RNAs by covariance-model searches with Infernal v1.1 against Rfam v15.1^32,33^.

Adapters and low-quality bases were removed using fastp^34^. Nuclear variants were called following the GATK v4.0 best-practices workflow^35^. Filtered reads were aligned to the *P. bianor* nuclear reference genome using BWA-MEM2 v2.2.1^36^. HaplotypeCaller was used to generate one genomic VCF file for each individual and chromosome, and these files were combined into chromosome-level cohort GVCFs using CombineGVCFs.

We constructed three nuclear datasets: dataset (a), a SNP-only dataset containing all samples; dataset (b), a full-site dataset containing variant and invariant sites for all samples; dataset (c) a SNP-only dataset containing only *P. bianor* samples for demographic history simulation. For datasets (a), SNP-only chromosome-level VCFs were generated using GenotypeGVCFs and merged across chromosomes. SNPs were extracted using SelectVariants and hard-filtered in GATK using the criteria QD < 2.0, MQ < 40.0, FS > 60.0, SOR > 3.0, MQRankSum < −12.5 or ReadPosRankSum < −8.0. Multiallelic records, records failing the GATK filters and sites with more than 20% missing data (F_MISSING > 0.2) were removed using bcftools. For dataset (b), chromosome-level VCFs retaining all genomic sites were generated using GenotypeGVCFs with the --all-sites option and merged across all 30 chromosomes using MergeVcfs. SNPs were extracted using SelectVariants and hard-filtered in GATK using the criteria QD < 1.5, MQ < 30.0, FS > 80.0, SOR > 4.0, MQRankSum < −15.0 or ReadPosRankSum < −10.0. Multiallelic records, sites failing the GATK filters, records containing the * alternative allele and sites with more than 20% missing data (F_MISSING > 0.2) were removed using bcftools. Invariant sites were extracted separately, retained when missingness was below 20% (F_MISSING <=0.2), adjusted for ploidy and filtered to remove records with missing alternative alleles. The filtered SNP and invariant-site datasets were subsequently merged using bcftools. For dataset (c), the filtering pipeline followed the same as dataset (a).

For mitochondrial genome reconstruction, filtered reads were aligned to the *P. bianor* mitochondrial reference using BWA-MEM2 v2.2.1. Alignments were converted to BAM format, coordinate-sorted, assigned read groups and processed using GATK to mark PCR duplicates. Deduplicated BAM files were indexed using SAMtools^37^. Mitochondrial variant and invariant sites were called using bcftools mpileup and call under a haploid model with ploidy set to 1, while retaining non-variant sites to reconstruct complete mitochondrial sequences.

To reduce interference from nuclear mitochondrial sequences, variant and invariant records were processed separately. Invariant sites were retained directly, whereas multiallelic variant records were temporarily decomposed into biallelic records using bcftools norm -m -any. Variant records were retained only when QUAL > 30, mapping quality > 30, total depth > 100 and the maximum alternative-allele depth was at least 10 reads in one or more samples. Records at the same genomic coordinate were then recombined using bcftools norm -m +any and concatenated with the retained invariant sites to produce the final mitochondrial VCF.

### Population genomic structure and phylogenetic reconstruction

Based on dataset (b), pairwise Weir and Cockerham’s *FST* values were estimated separately for the Z chromosome and the 29 autosomes using non-overlapping 20-kb windows implemented in pixy v2.0.0^38^. Genome-wide mean *FST* values were subsequently calculated for each population pair (Supplementary Table S2). Individual genome-wide inbreeding coefficients were estimated using the --het option in PLINK v1.9.0 based on autosomal part of dataset (b) only.

Population-structure analyses were based on dataset (a) from 319 ingroup individuals. Markers were pruned for linkage disequilibrium using an *r*² threshold of 0.2. Principal component analysis was conducted using VCF2PCACluster^39^, and individual ancestry proportions were estimated using ADMIXTURE v1.3.0^40^ for *K* = 1–12. The best-supported value of *K* was identified from the minimum cross-validation error (Supplementary Fig. 2).

Individual-level maximum-likelihood phylogenies were reconstructed using IQ-TREE v3.0.1^41^, with the outgroups included in all analyses. For the complete dataset of 325 individuals, we inferred a maximum-likelihood phylogeny using genome-wide common SNPs (MAF ≥ 0.05) with no missing sites from all autosomes. To examine cytonuclear phylogenetic discordance at a finer scale, we additionally selected 53 representative individuals spanning the major geographical populations (Supplementary Table S4) and independently reconstructed nuclear and mitochondrial phylogenies. The nuclear phylogeny was inferred from genome-wide autosomal SNPs retained after linkage-disequilibrium pruning at *r*² = 0.2, whereas the mitochondrial phylogeny was reconstructed from the complete mitochondrial-genome alignment. For each dataset, the best-fitting substitution model was selected using ModelFinder with -m MFP, and branch support was assessed with 1,000 ultrafast bootstrap replicates^42^. Ascertainment-bias correction (+ASC)^43^ was applied to analyses based on SNP datasets.

Evolutionarily significant units (ESUs) and finer geographical populations were defined using the ADMIXTURE results, the concatenated maximum-likelihood phylogeny and summaries of the window-based trees. Seven ESUs were recognized: *P. bianor*-Islands, *P. bianor*-Indochina, *P. bianor*-E-China, *P. bianor*-Himalaya, *P. dehaanii*, *P. okinawensis*-Okinawa and *P. okinawensis*-Amami. Introgression analyses were conducted independently at the ESU and geographical-population levels to distinguish gene flow among major evolutionary lineages from more localized exchange among regional populations.

### Genome-wide phylogenetic discordance and topology weighting analyses

Window-based phylogenomic analyses were conducted using the whole-genome SNP dataset described above, with a reduced set of representative individuals from each major taxonomic group (Supplementary Table S5) to improve computational efficiency. To minimize potential biases associated with coding regions, phylogenetic reconstruction was restricted to non-genic genomic regions. Gene coordinates were first sorted and merged using BEDTools v2.31.1 to generate continuous genic intervals. These regions were subsequently removed from chromosome coordinates, and the remaining intergenic regions were partitioned into non-overlapping 20-kb windows using bedtools makewindows^44^. Windows spanning fragmented intergenic regions were retained as generated.

Maximum-likelihood trees were independently inferred for each genomic window using IQ-TREE v3.0.1 with automatic model selection (-m MFP) and 1,000 ultrafast bootstrap replicates. Genome-wide gene-tree discordance was evaluated by comparing window trees with the dominant topology recovered from the individual-level maximum-likelihood phylogeny. Alternative topology frequencies and their distributions across the genome were summarized and visualized using TWISST^45^.

An autosomal species tree was reconstructed from the collection of window trees using ASTRAL-Hybrid^46^, which also estimated local quartet support and topology frequencies at internal nodes. PhyTop v0.3.2^47^ was subsequently applied to quantify the relative contributions of incomplete lineage sorting (ILS) and introgression (IH) to observed topological discordance based on the frequencies of alternative gene-tree topologies.

To further characterize lineage relationships across genomic compartments, topology weighting analyses were performed using TWISST^45^. For the three-species comparison involving *P. bianor*, *P. dehaanii*, and *P. okinawensis*, weights were calculated for all possible species-level topologies. Within *P. bianor*, topology weights were estimated among ESUs, with the Islands lineage further divided into LNY and YYM, and summarized for the five most frequent genome-wide topologies. Topology weights were subsequently calculated separately for autosomes and the Z chromosome to compare chromosome-specific patterns of phylogenetic relationships.

### Inference of introgression and incomplete lineage sorting patterns

Patterson’s D-statistics and *f*4-ratios were calculated using the Dtrios module in Dsuite v0.5^48^ (Supplementary Tables S6, S7). All triplets consistent with the corresponding guide tree were tested. Statistical significance was assessed by standard block jackknifing by dividing the dataset into 200 Jackknife blocks^49^, and combinations with *Z* > 3 were considered to show significant evidence of gene flow. Branch-specific introgression was further localized using the Dsuite f-branch framework based on the D-statistic results and the corresponding phylogeny, with *Z* > 3 again treated as evidence of significant introgression.

Potential hybrid origins were evaluated using HyDe^21^, with particular emphasis on the stable mixed ancestry observed in the E-China lineage. The analyses included four major *P. bianor* ESUs—Himalaya, Indochina, E-China, and Islands (Supplementary Table S10a). The run_hyde_mp.py module was initially used to exhaustively test possible parental combinations among geographical groups (Supplementary Tables S9a, S9b). Selected triplets of the form P1–Hybrid–P2 with *P* < 0.05 were subsequently evaluated using 100 bootstrap replicates. Mean parental contributions, expressed as γ, and their 95% bootstrap confidence intervals were calculated for each supported combination.

Gene flow among focal geographical populations was further evaluated using TreeMix v1.13. Several small populations were merged, and representative individuals were retained for analysis (Supplementary Table S11). The dataset was pruned for linkage disequilibrium at *R*² > 0.2 using PLINK v1.9.0-b.8 and converted to TreeMix format using plink2treemix.py. TreeMix first inferred a maximum-likelihood population tree and then tested models containing one to ten migration edges. Each model was repeated ten times to evaluate consistency. The optimal number of migration edges was determined using OptM v0.1.8^50^ with both the Evanno and Linear methods, and the selected one-migration model was visualized using plotting_funcs.R.

QuIBL^10^ was used to determine whether discordant gene trees were better explained by ILS alone or by ILS combined with introgression. Analyses focused on the ESUs used in Dsuite (Supplementary Table S8) and tested the same topology-guided triplets represented in the D-statistic and *f*4-ratio matrix. For each population-level triplet, all possible individual-level combinations were generated by selecting one individual from each unit. Window trees containing the three focal individuals and the designated outgroup were pruned to these four terminals while preserving branch lengths, and 1,000 retained trees were randomly sampled for each QuIBL run. Each individual-level triplet was treated as an independent replicate. For each of the three possible topologies, QuIBL estimated Bayesian information criterion values under an ILS-only model (BIC1) and a model incorporating both ILS and introgression (BIC2). We calculated ΔBIC as BIC2 − BIC1; following Feng et al.^51^, ΔBIC > 10 was interpreted as strong support for ILS alone, whereas ΔBIC < −10 was interpreted as strong support for ILS plus introgression. The ΔBIC values were converted into BIC-based model weights and combined with the relative frequency of the corresponding topology. ILS and introgression signals were first summarized within each individual triplet and then averaged across all individual combinations to obtain sample-balanced estimates for each population-level comparison.

### Bayesian multispecies coalescent analyses

BPP^52,53^ was used to estimate divergence times and test gene flow under the multispecies coalescent, which explicitly accounts for ancestral polymorphism, stochastic coalescence and incomplete lineage sorting (ILS).

The MSC-I model was applied to evaluate two categories of gene-flow hypotheses: (1) ancestral introgression scenarios suggested by TreeMix analyses, and (2) ancestral or recent gene flow between insular and continental lineages of *P. bianor*. For each tested lineage pair, reciprocal models were constructed to evaluate alternative directions of gene flow, except where unidirectional models were specified below. For the initial analyses, 1,500 CDS loci were randomly resampled before filtering. Complete CDS alignments, including both variant and invariant sites, were extracted from the VCF dataset. Loci shorter than 300 bp, longer than 10,000 bp, or containing fewer than 20 SNPs were excluded after random resampling.

For the first scenario, seven populations were represented by one individual each: Western, Central and Eastern Himalayan *P. bianor*; Izu *P. dehaanii*; Okinawa and Amami *P. okinawensis*; and the outgroup *P. paris* (Supplementary Table S12a). After filtering, 1,381 CDS loci remained. Twelve models tested reciprocal introgression between each of the three Himalayan lineages and either the ancestral branch of the two *P. okinawensis* populations or the common ancestral branch of *P. dehaanii* and *P. okinawensis*.

The second scenario comprised two initial sets of models. The first included 13 individuals from seven populations: two individuals each from Western Himalaya, Central Himalaya, Eastern Himalaya, Southwestern Thailand, Yaeyama and Lanyu, together with one *P. paris* outgroup (Supplementary Table S12a). After filtering, 1,386 CDS loci remained. Twenty-four models tested reciprocal gene flow between each of the four continental lineages and three insular configurations: the common ancestor of Yaeyama and Lanyu, extant Yaeyama and extant Lanyu.

The second set included 11 individuals from six populations: two individuals each from Southeastern Thailand, E-China, Taiwan, Yaeyama and Lanyu, together with one *P. paris* outgroup (Supplementary Table S12a). After filtering, 1,368 CDS loci remained. Eighteen models tested reciprocal gene flow between each of the three continental lineages and the three insular configurations described above. Additional models were constructed to specifically evaluate unidirectional introgression from the ancestral island lineage, extant Lanyu and extant Yaeyama into the common ancestor of the E-China and Taiwan lineages.

To assess the robustness of the inferred ghost introgression events to expanded population sampling, five additional MSC-I models were evaluated. These analyses included 11 sampled taxa, represented by one individual each: EG, EH, CH, WH, SET, SWT, EC, TW, YYM, LNY and the outgroup *P. paris* (Supplementary Table S12b). The five models specified unidirectional introgression from a ghost lineage associated with the ancestral Yaeyama–Lanyu lineage into one of five recipients: the common ancestor of EC and TW, WH, SWT, CH or SET. An additional randomly resampled CDS dataset was used for these analyses, with 1,300 CDS loci retained after applying the same filtering criteria for each model. The models involving the common ancestor of EC and TW, WH, and SWT are illustrated in Supplementary Fig. 12; complete results for all five models are provided in Supplementary Table S12b.

For all MSC-I analyses, the admixture proportion parameter (φ) was assigned a Beta(1,1) prior, and the population-size parameter (θ) was explicitly sampled under an inverse-gamma prior of IG(3, 0.014). The root-age parameter (τ) was assigned an IG(3, 0.15) prior in the initial analyses^13^ and an IG(3, 0.08) prior in the five analyses with expanded population sampling. Absolute divergence times were estimated by calibrating the root divergence between *P. bianor* and *P. paris* to approximately 10.57 Ma using a gamma approximation with shape = 64 and rate = 6.1^15^.

Chains for the first scenario were run for 200,000 iterations after a burn-in of 100,000 iterations, whereas those for the initial analyses of the second scenario were run for 400,000 iterations after a burn-in of 200,000 iterations. Samples were recorded at every iteration in these analyses. For each of the five models with expanded population sampling, chains were run for 1,800,000 iterations after a burn-in of 300,000 iterations, with samples recorded every three iterations to obtain 600,000 posterior samples. All other analytical procedures were the same as those used in the initial analyses. Statistical support for gene flow was assessed using Bayes factors approximated by the Savage–Dickey density ratio.

### Demographic history simulations

Historical changes in effective population size were reconstructed using PSMC v0.6.5^54^. Individuals with sequencing depth above 15× were retained, comprising 100 *P. bianor* samples. Multiple individuals per population were analysed to evaluate the consistency of inferred demographic trajectories. Diploid consensus sequences were generated from read alignments using bcftools and vcfutils.pl and converted to PSMC format using fq2psmcfa -q 20, which retained base calls with a minimum Phred-scaled quality score of 20. PSMC was run using the parameters -N25 -t10 -r5 -p “4+25*2+4+6”. Results were visualized using easy_psmc_plot.py, assuming a generation time of 0.3 years, equivalent to approximately three generations per year, and a mutation rate of 1.3 × 10⁻⁹ substitutions per site per generation^55^.

Alternative demographic scenarios were evaluated using fastsimcoal2 v2.8 under a composite-likelihood framework based on simulated site-frequency spectra^56^. We investigated demographic relationships among the four major *P. bianor* ESUs: Himalaya, Indochina, E-China and Islands (Supplementary Table S13). The analysis was conducted using dataset (c), in which variants were first filtered to retain only complete sites without missing genotypes and subsequently pruned for linkage disequilibrium at *R*² > 0.2 using PLINK v1.9.0 to minimize biases caused by correlated markers^57^. Two-dimensional folded site-frequency spectra (2D-SFS) were generated using easySFS.

Eight alternative demographic models were compared for the four-lineage system, including two strictly bifurcating models and six hybrid-origin models describing alternative scenarios for the formation of the E-China lineage (Supplementary Fig. 13). All hybrid-origin models incorporated gene flow between E-China and other major lineages, but differed in the timing and evolutionary context of admixture events. In particular, model PB6 represented simultaneous divergence and admixture between E-China and the Islands lineage; model PB7 assumed that E-China first diverged as an independent lineage followed by subsequent introgression from the Islands lineage; and model PB8 incorporated a scenario in which an unsampled island-related ghost lineage diverged concurrently with E-China and contributed genetic material through later introgression. Continuous migration after lineage formation was additionally modeled between E-China and the other major lineages.

Each demographic model was evaluated using 50 independent optimization replicates, with 100,000 coalescent simulations per optimization cycle and 50 optimization cycles. Analyses assumed three generations per year and a mutation rate of 1.3 × 10⁻⁹ substitutions per site per generation. Models were ranked using the Akaike information criterion (AIC; Supplementary Fig. 13). Parameter uncertainty for the best-supported model was assessed using 100 parametric-bootstrap site-frequency-spectrum replicates generated from maximum-likelihood parameter estimates. Mean parameter estimates and 95% confidence intervals were calculated following the fastsimcoal2 manual.

### Ecological niche modelling under present-day and palaeoclimatic scenarios

Species distribution models (SDMs) were calibrated separately for Himalayan and Eastern China lineages using present-day occurrence records and environmental predictors, and projected onto Last Glacial Maximum (LGM; ca. 21 ka) climatic conditions. Models were constructed using a presence-background maximum entropy framework^58,59^. Occurrence records were filtered for geographic validity, spatially thinned at 10-km intervals using an equal-area projection (EPSG:6933), and restricted to one record per 2.5-arc-min environmental grid cell. Lineages with fewer than 15 environmentally complete occurrences were excluded. The accessible calibration area (*M*) was defined as a 600-km buffer around retained occurrences following the accessible-area concept in ecological niche modelling^60^. Up to 10,000 background cells were randomly sampled within *M*, excluding cells within 10 km of occurrence localities.

Environmental predictors were obtained from PaleoClim bioclimatic layers^61^, including BIO1, BIO4, BIO6, BIO12, BIO15, BIO18 and BIO19. Variables with zero variance were removed, and highly correlated predictors (|r| > 0.80) were pruned based on an ecological prioritization scheme favoring climatic extremes and seasonal variability. Variables were retained according to the following priority: BIO6, BIO4, BIO15, BIO18, BIO19, BIO1 and BIO12. This prioritization emphasizes climatic factors associated with physiological constraints and seasonal niche differentiation in ectotherms while minimizing redundancy among correlated climatic variables. A maximum of five predictors was retained according to sample size.

Models were fitted using maxnet implemented in ENMeval v2.0.5^62,63^. Linear (L) and linear–quadratic (LQ) feature classes were evaluated with regularization multipliers of 1, 2 and 3. Model performance was assessed using geographically structured spatial cross-validation with up to three folds (minimum five occurrences per fold) to reduce inflation of predictive performance caused by spatial autocorrelation^64^. Models were first filtered using an omission-rate threshold (OR10 ≤ 0.10) and subsequently ranked by AICc and model complexity. When no model met this criterion, models within 0.02 of the minimum OR10 were considered.

Present-day and LGM climatic layers were obtained from PaleoClim at 2.5-arc-min resolution and standardized to a common spatial grid before projection. LGM projections represent climatic suitability under glacial conditions without additional palaeoshoreline reconstruction or sea-level correction. Elevation was excluded because applying modern topography to palaeoclimatic scenarios may introduce unrealistic constraints, particularly in regions affected by historical landscape changes.

Optimized models were projected onto current and LGM environmental conditions using corresponding climatic predictors. Habitat suitability was expressed on the cloglog scale (0–1), and binary suitable areas were defined using the 10-percentile training-presence threshold. Predictor importance was evaluated using permutation-based AUC decreases, response curves were generated for retained predictors, and extrapolation risk was assessed by identifying predictors exceeding calibration ranges. Final projections were restricted to 60–130°E and 5–45°N and displayed using a common suitability scale.

## Declarations

### Ethics approval and consent to participate

All procedures involved in the tissue collection of animals were in accordance with the China Practice for the Care and Use of Laboratory Animals and approved by the China Zoological Society (permit number: GB/T35892-2018).

### Consent for publication

Not applicable.

### Availability of data and materials

The newly reported WGS data will be made publicly available upon formal publication of this article in a peer-reviewed journal.

Public WGS data were obtained from PRJNA892033, PRJNA1009405 and PRJNA1043105. Transcriptomic reads and orthologous protein sequences used for genome annotation were obtained from PRJNA1009405, respectively. Sample-level provenance and supporting results are provided in the supplementary files.

## Authors’ contributions

HW and YC contributed to conceptualization, HW conducted sample collection, analyses and visualization. HW, TW and BX contributed to methodology and drafting. YC acquired funding. HW, YC and BX reviewed and edited the manuscript.

## Acknowledgements

The work was supported by the National Key Research and Development Program of China (2023YFF1304800 and 2024YFF1307500), the International Partnership Program of Chinese Academy of Sciences for Grand Challenges (073GJHZ2023091GC). We thank D. Lin of the University of Tokyo, Japan; Z. Zhou of Yunnan Province, China; Mr. Xie of Anhui Province, China; Z. Wang of Dankook University, South Korea and others for their assistance with sample collection. We thank J. Liu of Liaoning Province, China for assistance with the literature search. We are grateful to Prof. Xiaosheng Chen of College of Forestry and Landscape Architecture, South China Agricultural University for providing laboratory equipment during the exploratory phase of this project, and to Prof. John S. Ascher of the Department of Biological Science, National University of Singapore for his valuable advice during this phase. We also thank Qi Xiao and labmates at the Chengdu Institute of Biology for their helpful advice on data analyses.

