## Supplementary Materials for "Deep ancestral structure, admixture and peripheral persistence shape diversity in the *Papilio bianor* species complex"

### Supplementary Table

All supplementary tables are available in the attached XLSX file (Supplementary Table).

### Supplementary Figure

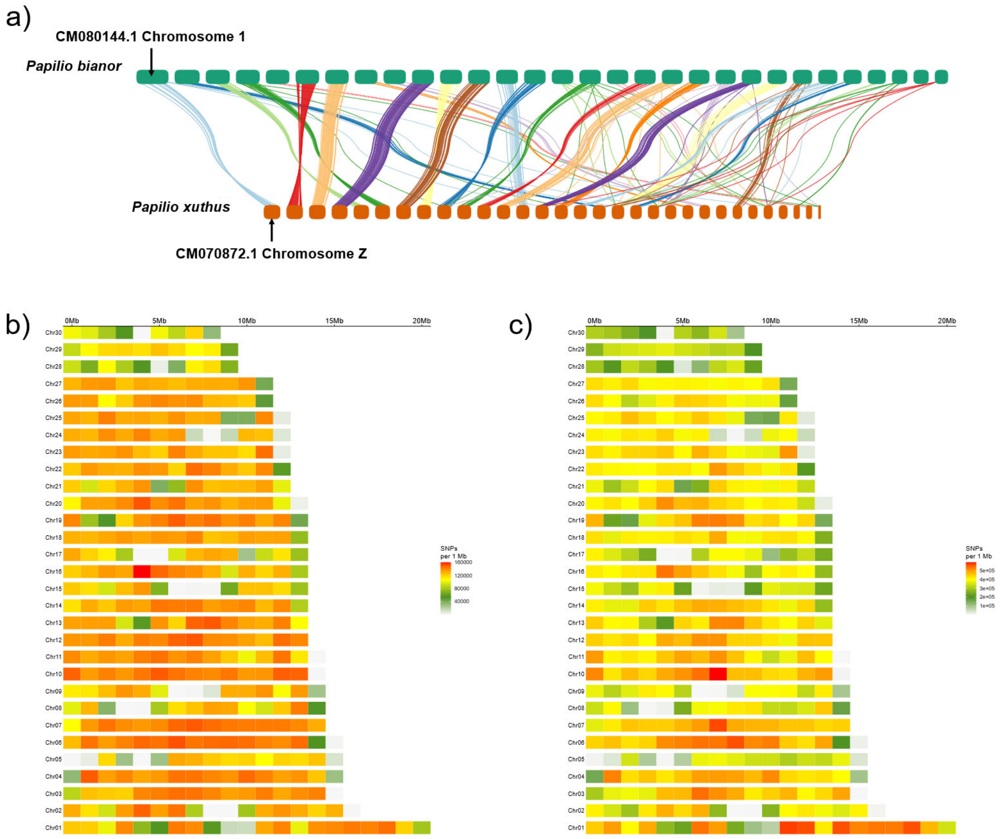


**Supplementary Fig. 1. Chromosomal synteny and genome-wide distribution of single-nucleotide variant density.** (a) Chromosomal synteny analysis between *Papilio bianor* and *P. xuthus*, confirming Chr01 as the Z chromosome of *P. bianor*. (b) Genome-wide density distribution of the 47,828,665 SNPs retained after stringent filtering of dataset (a), and the 49,039,202 SNPs of dataset (b) (invariant sites excluded); comprising 325 individuals including the outgroups, across all 30 chromosomes.

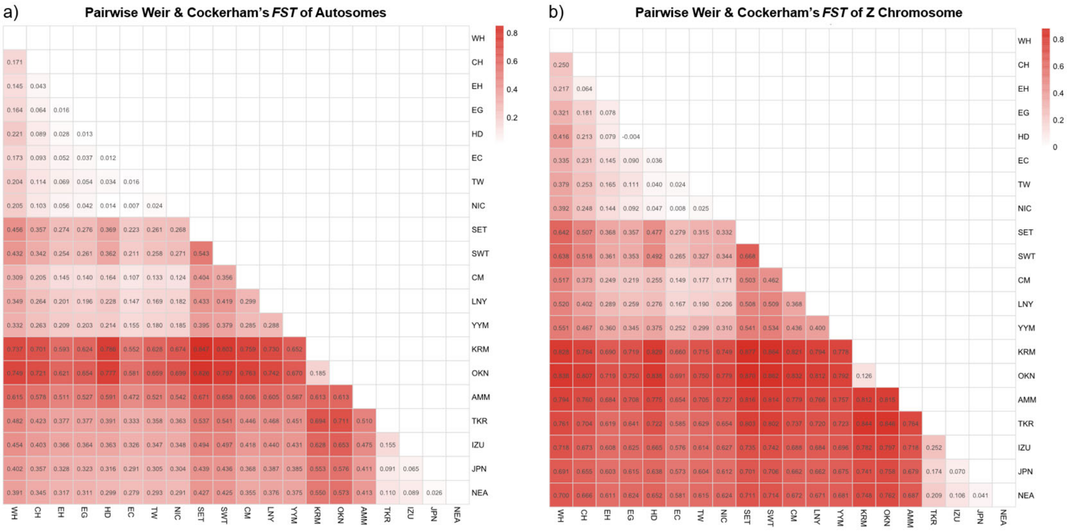


**Supplementary Fig. 2. Genome-wide average pairwise Weir and Cockerham's $F_{ST}$ among geographical populations.** Pairwise $F_{ST}$ values were estimated separately for the autosomes (left) and Z chromosome (right). Numbers within cells indicate genome-wide mean $F_{ST}$, and colour intensity represents the magnitude of genetic differentiation, with darker red indicating greater differentiation. Population abbreviations (Supplementary Table S1b) and the samples included in each population (Supplementary Table S2) are provided in Supplementary Tables 1 and 2, respectively.

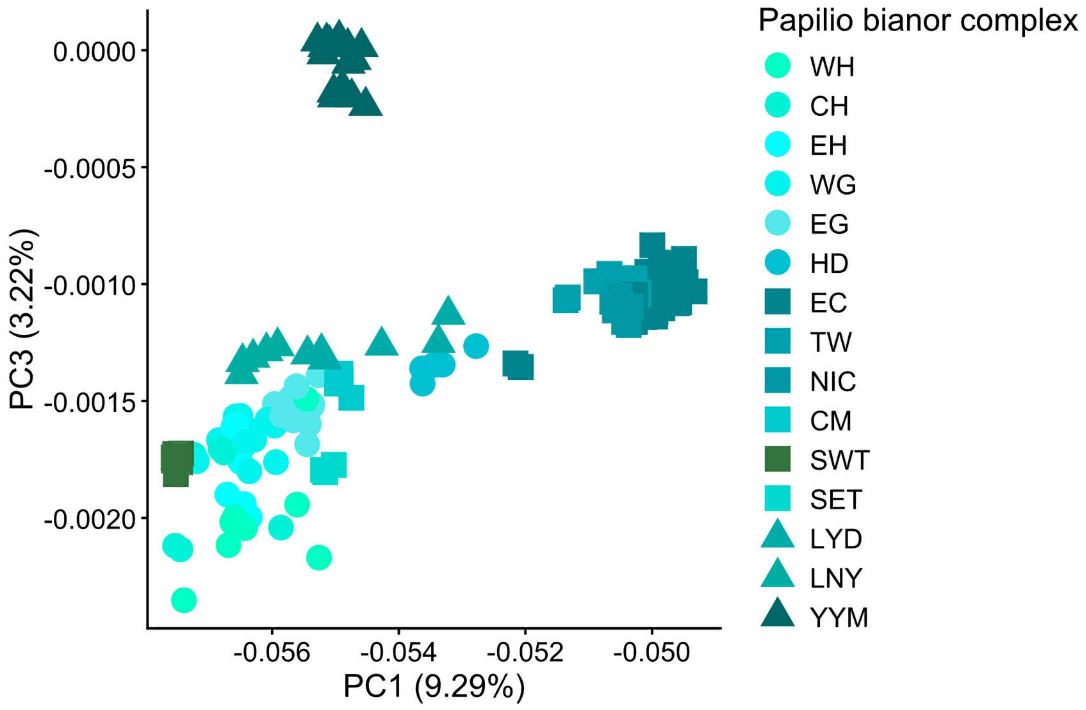


**Supplementary Fig. 3. Principal component analysis of *Papilio bianor* populations.** Principal component plots showing the relationships between PC3 and PC1 within *P. bianor*.

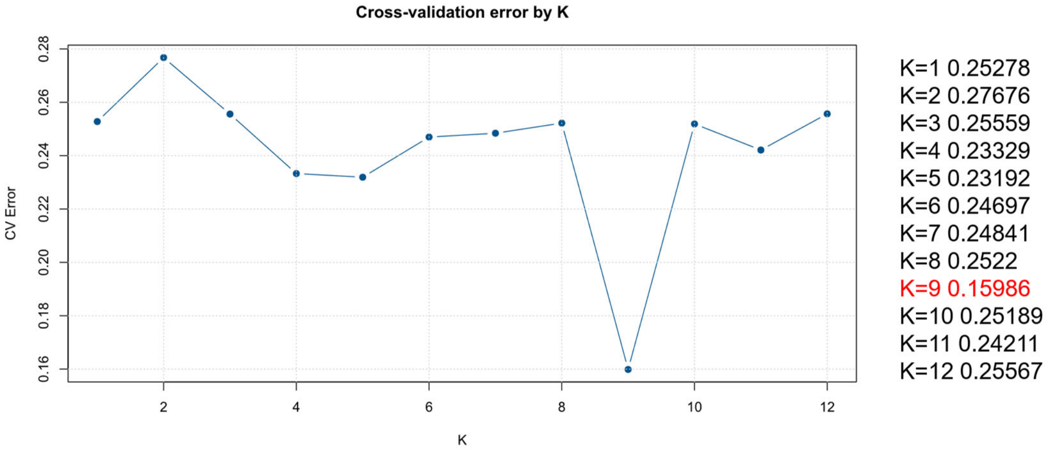


**Supplementary Fig. 4. Cross-validation error values of ADMIXTURE analyses across different numbers of ancestral clusters (K = 1–12).** The cross-validation (CV) error was used to evaluate model performance, with lower values indicating better model fit. The CV error reached its minimum significantly at K = 9, indicating the optimal number of ancestral components.

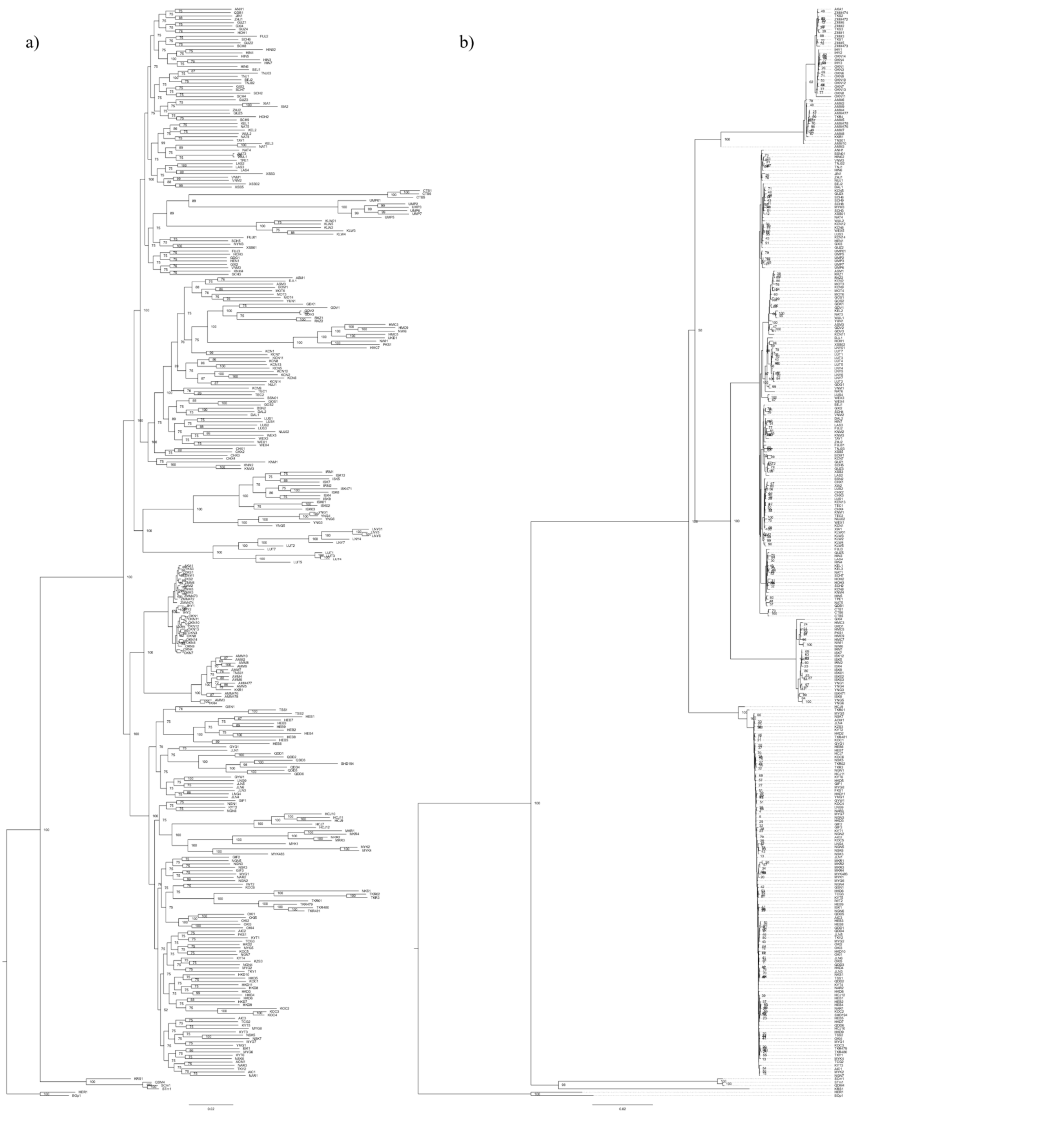


**Supplementary Fig. 5. Maximum-likelihood phylogenetic trees inferred from nuclear and mitochondrial genomes for all 325 samples (Supplementary Table S1a).** (a) The nuclear genome phylogeny was reconstructed using genome-wide common variants (MAF ≥ 0.05) under the best-fit nucleotide substitution model selected by BIC (GTR+F+R10). (b) The mitochondrial genome phylogeny was inferred from a 15,333-bp mitochondrial sequence alignment under the best-fit model selected by BIC (GTR+F+R5). Bootstrap support values from 1,000 replicates are shown for all nodes in both phylogenetic trees.

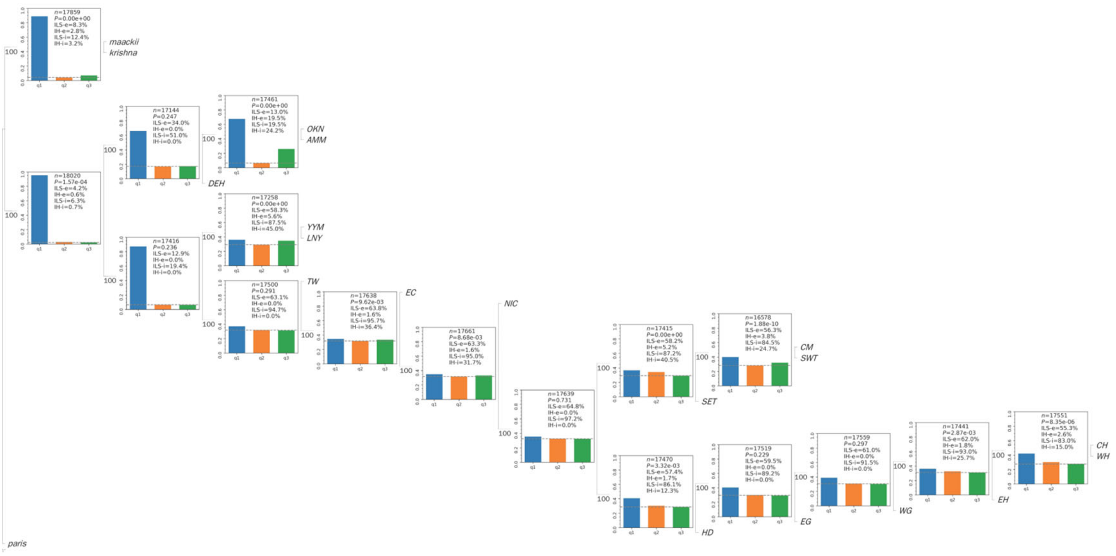


**Supplementary Fig. 6. ASTRAL-Hybrid species tree and node-specific topological discordance inferred from autosomal window trees.** Numbers beside branches indicate local posterior probabilities. At each internal node, PhyTop bar plots show the frequencies of the dominant topology (q1, blue) and two alternative topologies (q2, orange; q3, green). The dashed line represents the expected frequency of each alternative topology under incomplete lineage sorting (ILS) alone. *n* is the number of informative window trees, and *P* is the chi-square-test probability for equal frequencies of q2 and q3. ILS-e and IH-e denote the proportions of topologically discordant window trees attributed to ILS and introgression/hybridization (IH), respectively, whereas ILS-i and IH-i quantify their estimated strengths. Within *P. bianor*, ILS-i ranged from 83.0% to 97.2%, while non-zero IH-i values ranged from 12.3% to 45.0%.

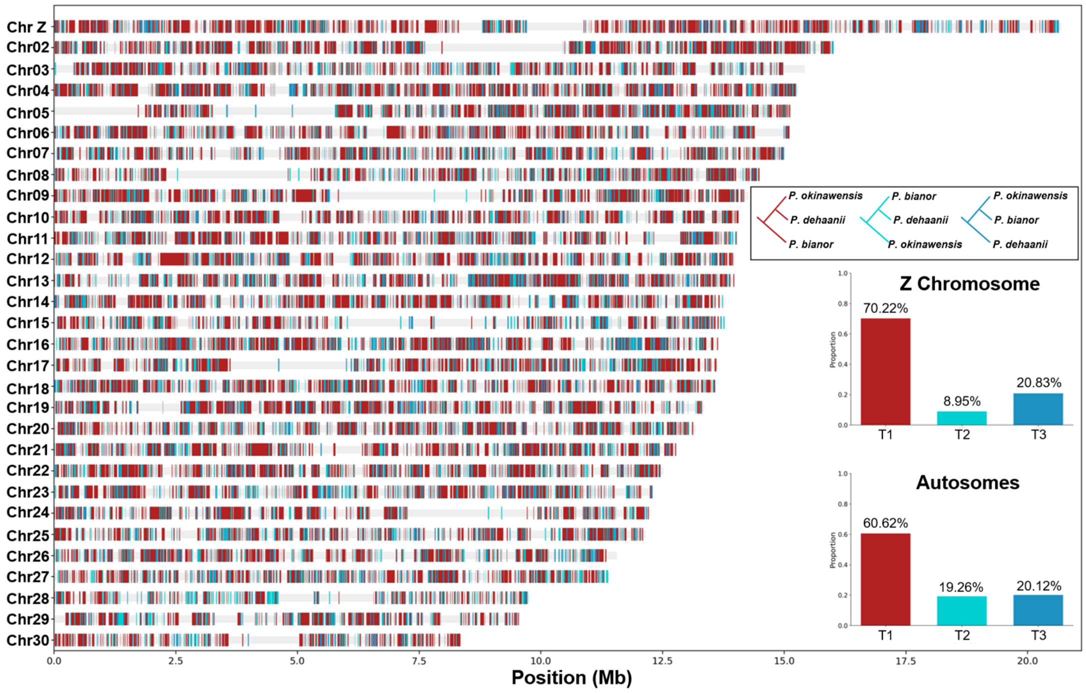


**Supplementary Fig. 7. Genome-wide distribution of three alternative topologies among *Papilio bianor*, *P. dehaanii*, and *P. okinawensis* inferred using Twisst.** Coloured segments show the distribution of local topologies along the Z chromosome and autosomes, with genomic positions given in megabases. T1 (red) represents (*P. bianor*, (*P. okinawensis*, *P. dehaanii*)); T2 (cyan) represents (*P. okinawensis*, (*P. bianor*, *P. dehaanii*)); and T3 (blue) represents (*P. dehaanii*, (*P. okinawensis*, *P. bianor*)). The bar plots show the proportions of the three topologies on the Z chromosome and autosomes. T1 accounted for 70.22% of the Z chromosome and 60.62% of the autosomes; T2 accounted for 8.95% and 19.26%, respectively; and T3 accounted for 20.83% and 20.12%, respectively.

15:36

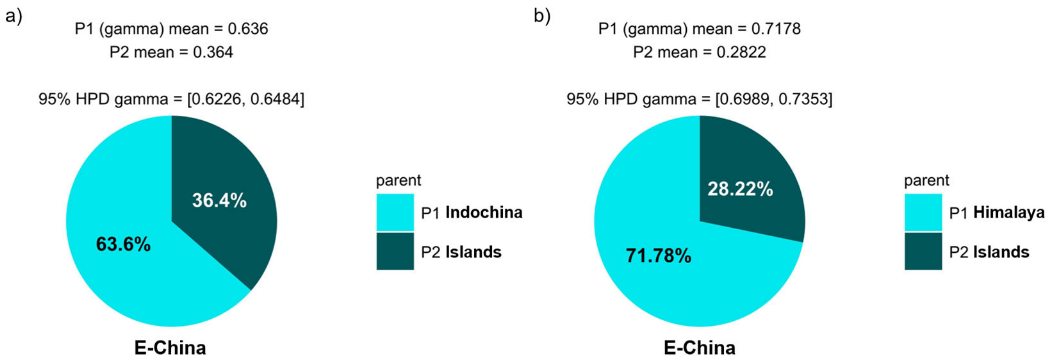


**Supplementary Fig. 8. Detection of hybrid origin signals and bootstrap estimation of ancestral contributions for the *Papilio bianor*-Eastern China lineage using HyDe.** Hybridization tests were conducted under the hybrid-origin hypothesis, with cyan indicating the contribution from mainland lineages and dark green indicating the contribution from island lineages. Pie charts show the mean estimated parental contributions (γ) and their corresponding 95% highest posterior density (HPD) intervals based on 100 bootstrap replicates (Supplementary Tables S10b, S10c). The sample composition and population assignments (Supplementary Table S10a) are provided in Supplementary Table S8, and the results of all tested triplets (Supplementary Table S9b) are summarized in Supplementary Table S9.

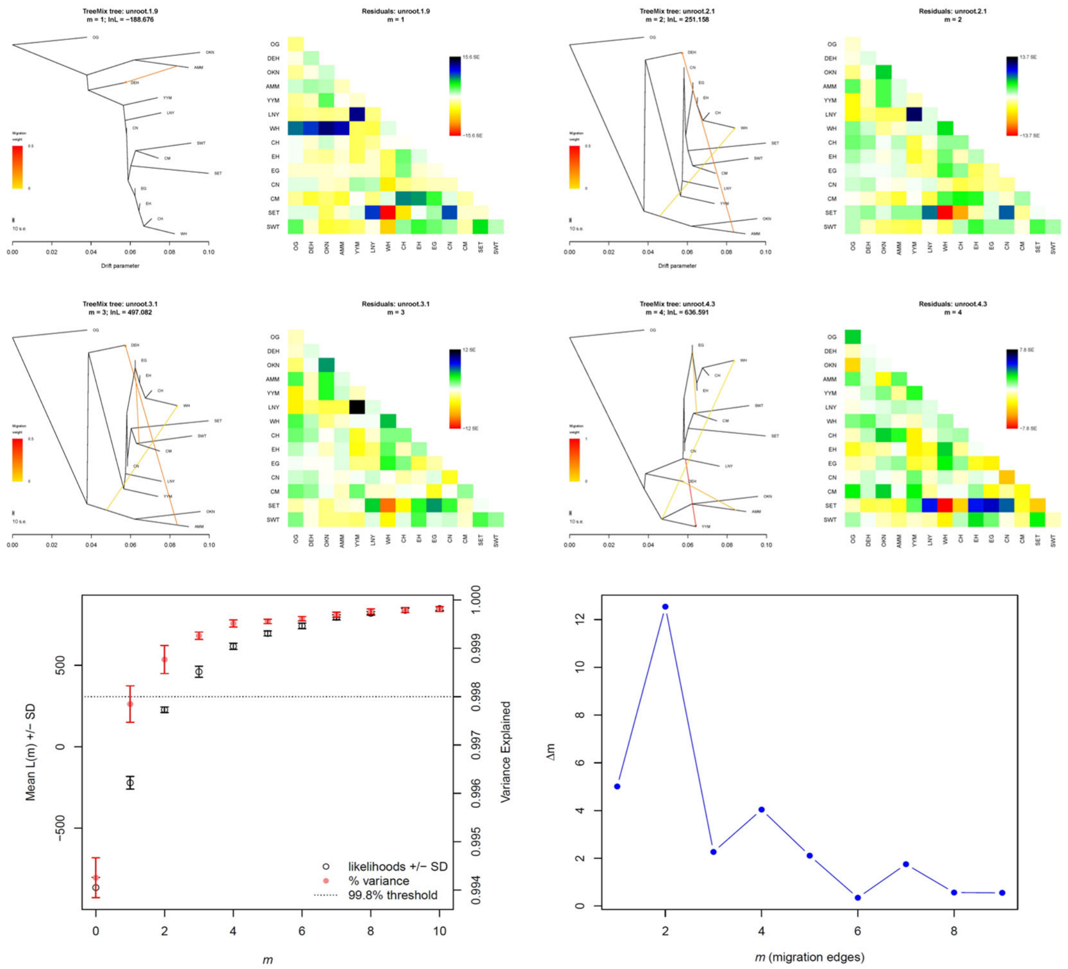


**Supplementary Fig. 9. TreeMix analysis of population relationships and migration among *Papilio* lineages.** Models with 1-10 migration edges were evaluated, and the maximum-likelihood population graphs and residual covariance matrices are shown for m=1-4. Horizontal branch lengths are proportional to genetic drift. Migration edges are coloured from yellow to red, with redder colours indicating larger migration weights. Residuals are expressed in standard-error units; positive residuals indicate greater observed covariance than predicted by the model, whereas negative residuals indicate lower observed covariance. The lower-left panel shows the mean log-likelihood ± SD and the proportion of variance explained for models with m=1-10; the dotted line marks the 99.8% variance-explained threshold. The lower-right panel shows the OptM Δm statistic. Both the variance-explained threshold and the maximum Δm identified m=2 as the optimal number of migration edges.

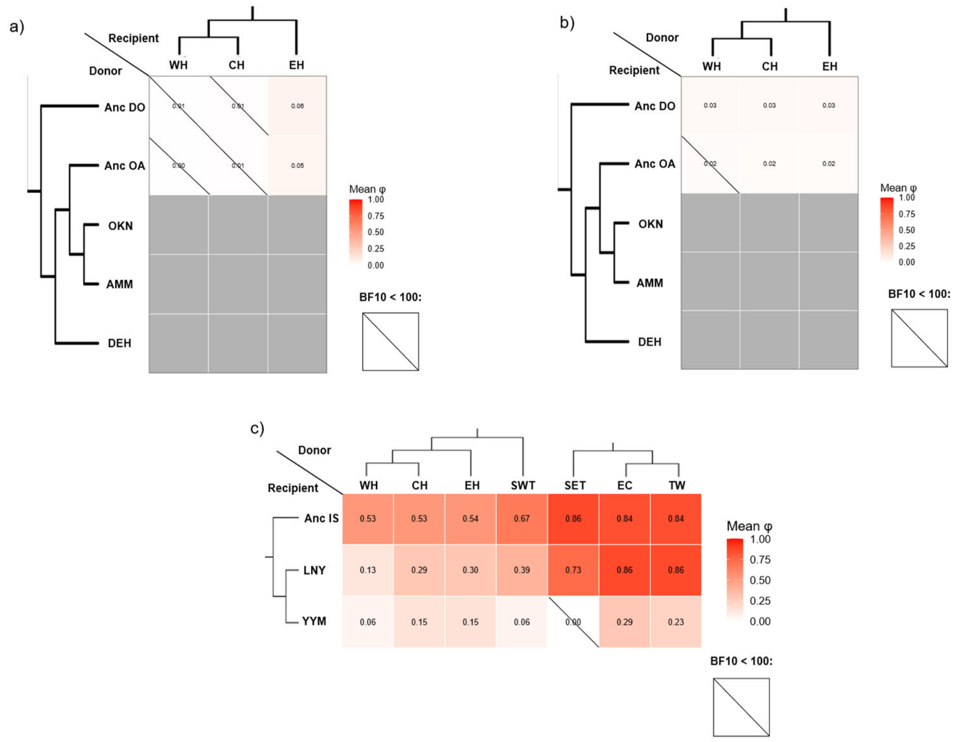


**Supplementary Fig. 10. Directional introgression scenarios tested using BPP under the multispecies coalescent with introgression (MSC-I) model.** Heatmaps show the posterior mean introgression proportion (φ) for each tested donor–recipient pair (Supplementary Table S12b), with donor and recipient axes indicated separately in each panel. Grey cells represent combinations that were not tested. (a) Introgression from the common ancestor of *P. dehaanii* and *P. okinawensis* (Anc DO) or the ancestral *P. okinawensis* lineage (Anc OA) into the western (WH), central (CH), and eastern (EH) Himalayan lineages of *P. bianor*. (b) Reciprocal scenarios in which WH, CH, and EH were treated as donors and Anc DO or Anc OA as recipients. (c) Introgression from the ancestral insular lineage shared by the Yaeyama Islands and Lanyu lineages (Anc IS), the extant Lanyu lineage (LNY), or the extant Yaeyama Islands lineage (YYM) into seven continental or near-continental *P. bianor* lineages: WH, CH, EH, southwestern Thailand (SWT), southeastern Thailand (SET), eastern China (EC), and Taiwan (TW). Cell values and colour intensity represent posterior mean φ, with darker red indicating larger estimates. Diagonal slashes identify scenarios with $BF_{10} < 100$, whereas cells without slashes met the criterion for strong support ($BF_{10} \geq 100$). Bayes factors were approximated using the Savage–Dickey density ratio. Detailed results are provided in Supplementary Table S10.

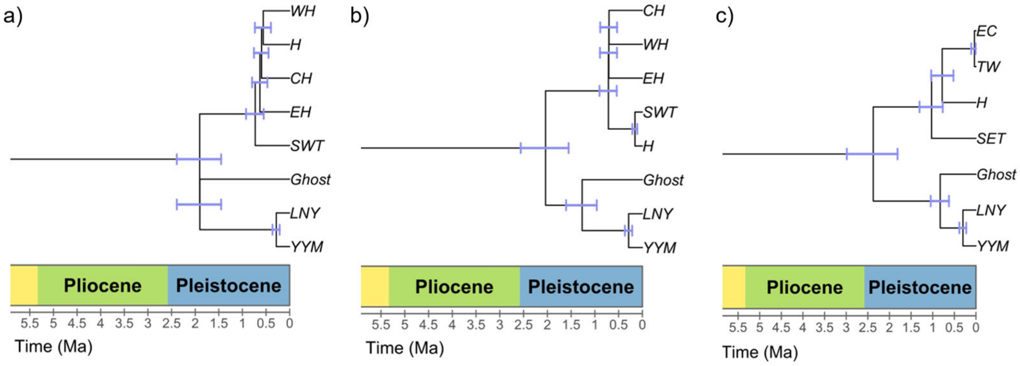


**Supplementary Fig. 11. Root-calibrated BPP MSC-I chronograms for three scenarios of ancient introgression involving the ancestral island lineage.** The three models place the lineage associated with the introgression event (H) adjacent to **(a)** WH, **(b)** SWT, or **(c)** the EC–TW lineage. The estimated ingroup root ages were 1.91 Ma (95% HPD: 1.45–2.39 Ma), 2.05 Ma (1.55–2.56 Ma), and 2.39 Ma (1.81–2.99 Ma), respectively. In panel **(a)**, the principal continental divergence times were 0.73 Ma (0.55–0.92 Ma), 0.63 Ma (0.47–0.80 Ma), 0.60 Ma (0.45–0.76 Ma), and 0.56 Ma (0.40–0.73 Ma). In panel **(b)**, the corresponding estimates were 0.72 Ma (0.54–0.90 Ma), 0.71 Ma (0.54–0.89 Ma), 0.70 Ma (0.53–0.89 Ma), and 0.16 Ma (0.11–0.21 Ma). In panel **(c)**, SET diverged from the remaining continental lineages at 1.03 Ma (0.77–1.30 Ma), H diverged from the EC–TW lineage at 0.78 Ma (0.52–1.04 Ma), and EC and TW diverged at 0.045 Ma (0.007–0.109 Ma). The divergence between LNY and YYM was estimated at 0.285 Ma (0.211–0.363 Ma), 0.291 Ma (0.215–0.369 Ma), and 0.303 Ma (0.225–0.384 Ma) in panels **(a–c)**, respectively. Blue bars denote 95% HPD intervals, and the time scales are shown in millions of years ago (Ma). Ghost denotes the unsampled ancestral lineage included in each MSC-I model.

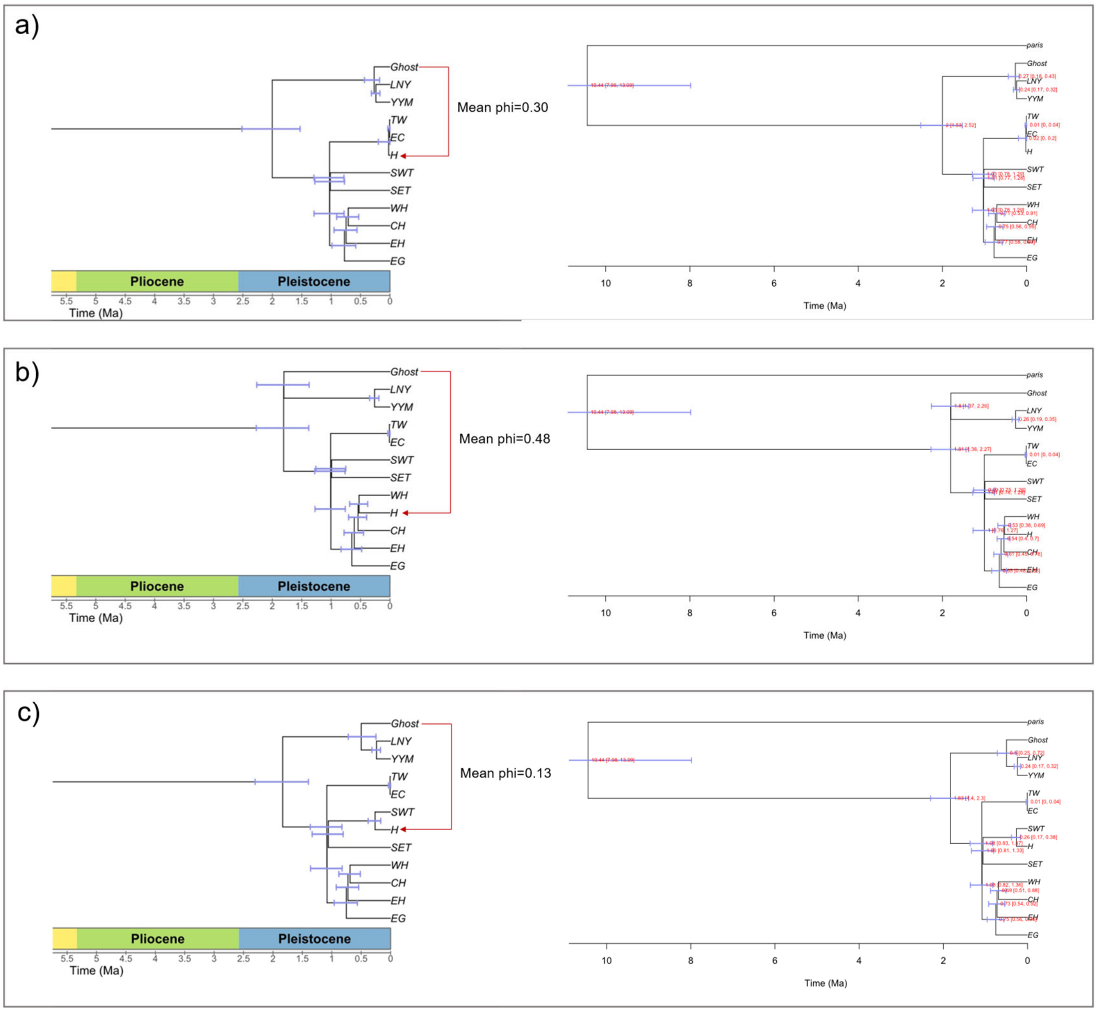


**Supplementary Fig. 12. BPP MSC-I replication analyses of ghost introgression under expanded population sampling.** Each row presents two visualizations of the same MSC-I model: the left panel shows the inferred introgression network and posterior mean introgression probability (φ), whereas the right panel shows the time-calibrated network with posterior median node ages and 95% highest posterior density intervals (blue bars; Ma). Red arrows indicate the direction of introgression, and *Ghost* denotes the inferred unsampled ancestral lineage. (a) Introgression into the common ancestor of EC and TW (mean φ = 0.30). Compared with the previous analysis based on limited population sampling, both the origin of the ghost lineage and the introgression event were inferred to be more recent, with a smaller ancestry contribution from the island lineage. (b) Introgression into WH (mean φ = 0.48). The inferred origin time of the ghost lineage, timing of introgression, and introgression proportion were broadly consistent with the previous model. (c) Introgression into SWT (mean φ = 0.13). Divergence-time estimates remained broadly consistent after additional populations were included, whereas the inferred contribution from the island lineage was substantially reduced. Complete results for all BPP MSC-I models under expanded population sampling are provided in Supplementary Table S12b.

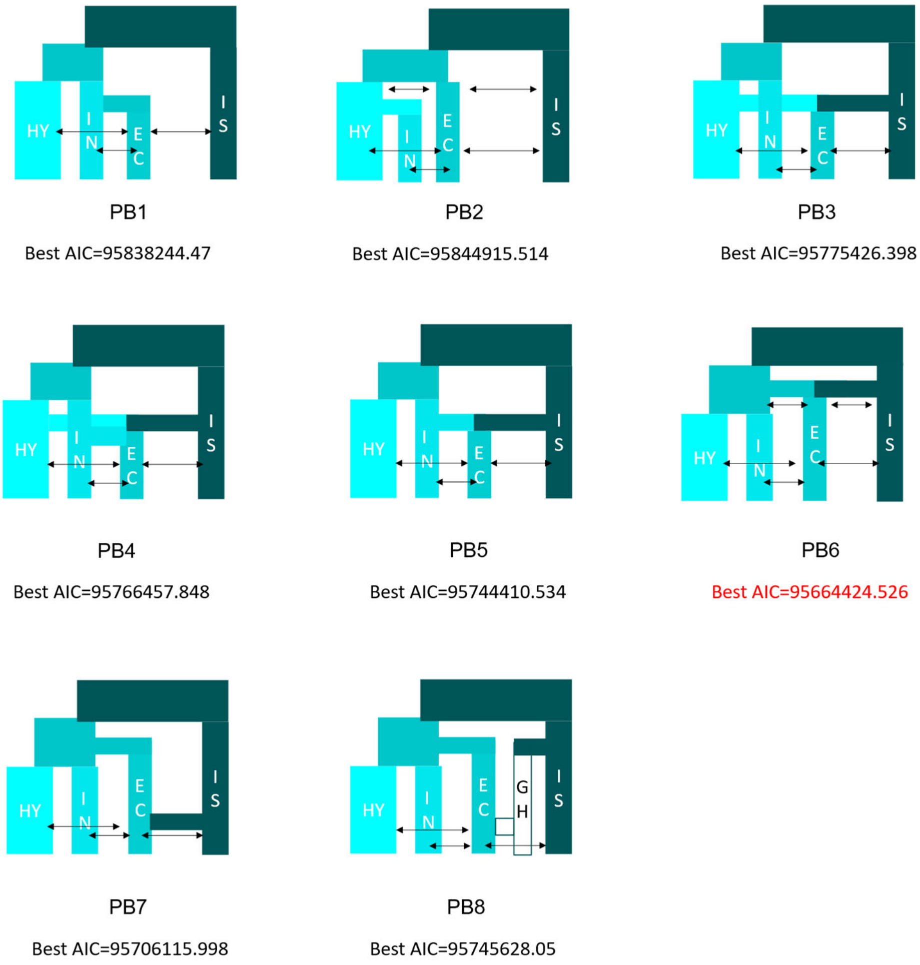


**Supplementary Fig. 13. Eight alternative demographic models tested in fastsimcoal2 for four evolutionarily significant units (ESUs) of *Papilio bianor*.** HY, Himalaya; IN, Indochina; EC, Eastern China; IS, Islands; GH, unsampled ghost lineage. PB1 and PB2 are bifurcating-divergence models with continuous migration, with EC–IN and HY–IN as sister lineages, respectively. PB3–PB5 model the hybrid origin of EC through admixture between HY and IS (PB3), among HY, IN, and IS (PB4), or between IN and IS (PB5). PB6 models EC as originating through admixture between the ancestral HY–IN population and IS. PB7 is a variant of PB6 with the post-admixture migration pattern shown. PB8 incorporates GH as an unsampled island-related contributor to EC. Double-headed arrows indicate continuous bidirectional migration. The lowest Akaike information criterion value across 100 independent replicates is reported for each model. PB6 had the lowest AIC (95,664,424.526) and was identified as the best-fitting model.

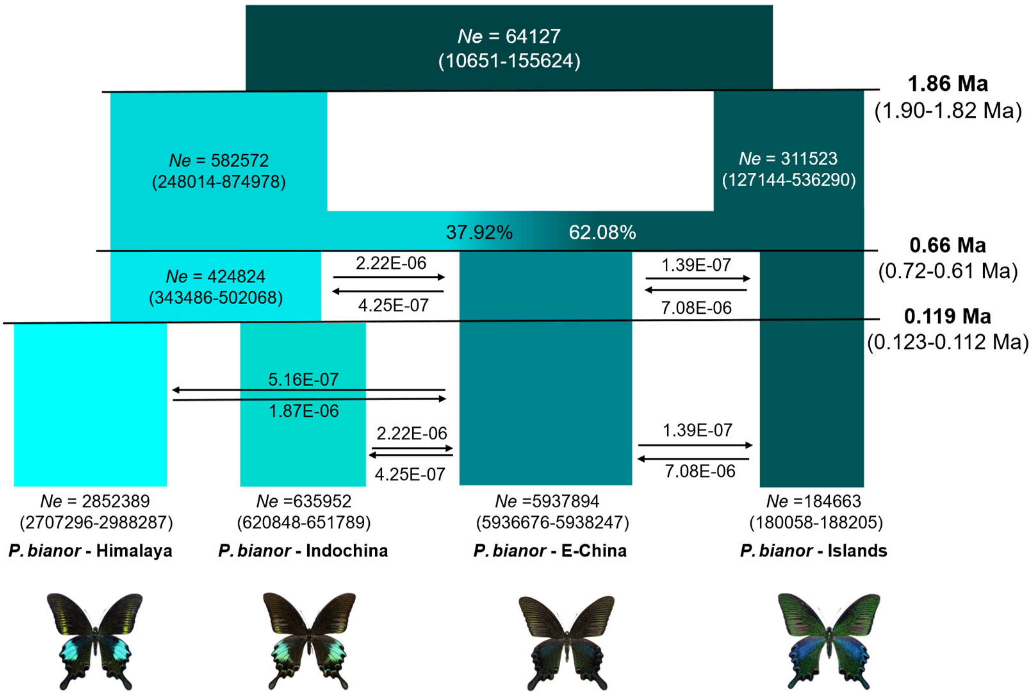


**Supplementary Fig. 14. Bootstrap parameter estimates for the best-fitting fastsimcoal2 demographic model (PB6) of four *Papilio bianor* ESUs.** The Islands lineage diverged from the ancestral mainland lineage at 1.86 Ma (95% HPD: 1.82–1.90 Ma). The Eastern China lineage originated at 0.66 Ma (0.61–0.72 Ma), with 37.92% ancestry from the ancestral Himalaya–Indochina population and 62.08% from the Islands lineage. The Himalaya and Indochina lineages diverged at 0.119 Ma (0.112–0.123 Ma). Effective population sizes ($N_e$) are shown for ancestral and extant populations. Horizontal arrows indicate the direction of continuous migration, with the corresponding migration rates shown above or below each arrow. Parameter values are bootstrap means, and values in parentheses are 95% HPD intervals (Supplementary Table S13).

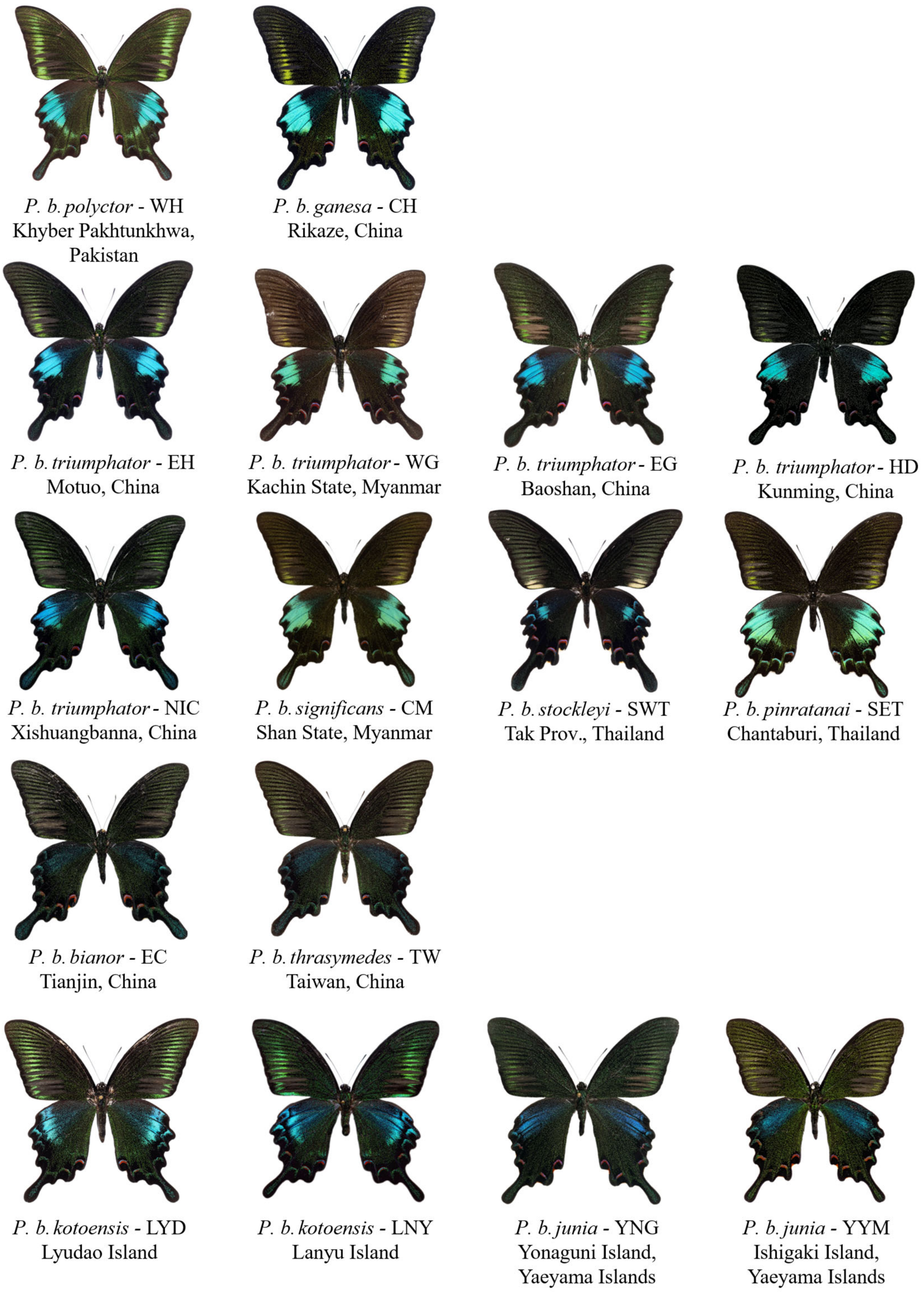


**Supplementary Fig. 15 Subspecies-level classification of geographic populations and representative specimen wing patterns.** Geographic groupings correspond to Supplementary Table 1, with the collection locality provided for each specimen. Specimen images illustrate wing-pattern characteristics only and do not represent actual relative sizes.

### Supplementary Discussion

#### Taxonomic implications

The *Papilio bianor* complex has long been taxonomically unstable. Continental forms, particularly the bright-green western Himalayan *polyctor* and darker eastern *bianor*, have repeatedly been treated as separate species, whereas *dehaanii* and *okinawensis* have often been classified as geographical subspecies of *P. bianor* sensu lato[65,66]. Recent systematic and hybridization studies increasingly support independent species status for *P. dehaanii* and *P. okinawensis*[17,67,68], but the status of western Himalayan *polyctor*, Yaeyama *junia* and other geographical forms of *P. bianor* remains unresolved[69,70]. Although some recent studies have restored *polyctor* as a species, they did not fully incorporate *junia*. Earlier multilocus analyses instead placed *polyctor* in a deeply divergent clade with *junia* and a Chinese sample, rather than as the sister lineage to all remaining continental *P. bianor*[15]. Our genomic results place these contrasting classifications within a history shaped by ancestral polymorphism, ancient admixture and subsequent regional differentiation.

The broad geographical and phenotypic variation of *P. bianor* also makes it informative for understanding diversification across the Oriental region. Himalayan wing-pattern variation broadly corresponds to the Western, Central and Eastern Himalayan climatic regions, represented by *polyctor*, *ganesa* and *triumphator*, respectively[20,65,71–73]. In Indochina, geographically restricted populations from the Shan Plateau, Tenasserim Range and Cardamom Mountains exhibit pronounced differences in wing shape and coloration, although morphologically intermediate populations occur in contact zones[70,74]. Eastern Chinese, Taiwanese, Lanyu–Lyudao and Yaeyama populations similarly show combinations of regional differentiation and morphological transition[65,75]. The close correspondence between phenotypic variation and major biogeographical regions suggests roles for both geographical isolation and local environmental adaptation, whereas narrow transition zones indicate that gene flow has not completely erased lineage differentiation.

Our results identify four principal evolutionarily significant units (ESUs) within *P. bianor*: Himalaya, Indochina, E-China and Islands. The deepest genome-wide division separates the Islands ESU, comprising *P. b. junia* from the Yaeyama Islands and *P. b. kotoensis* from Lanyu and Lyudao, from the mainland radiation. Although these two major lineages diverged approximately two million years ago, their subsequent histories remained connected through episodes of admixture. The strongly supported admixed origin of the Eastern China lineage and island-related ancestry in continental populations demonstrate that deep divergence and historical genetic exchange have jointly shaped present-day diversity. Likewise, the high differentiation of some peripheral populations, with FST values comparable to or exceeding those in some interspecific comparisons, may reflect the combined effects of ancient ancestry mixing, geographical isolation and genetic drift.

Within the mainland radiation, gene flow among the Himalayan, Indochinese and E-China ESUs was generally weak, and the observed hybrid zones showed no indication of expansion. The persistence of distinct genomic units despite localized contact suggests that lineage boundaries are being maintained and that speciation may be underway. Pronounced population structure also occurs within ESUs, particularly in peripheral Himalayan and Indochinese populations, indicating that differentiation is proceeding at multiple hierarchical levels. These patterns identify both the major ESUs and their differentiated constituent populations as important units for evolutionary investigation and conservation.

The present results nevertheless do not provide a sufficiently reliable basis for dividing the sampled *P. bianor* lineages into separate species. Divergence time, elevated FST, mitochondrial distinctiveness and diagnostic morphology document evolutionary differentiation, but their taxonomic interpretation must account for the reticulate history of the complex. Formal species-level revision should therefore be grounded in clear evidence of reproductive isolation, integrated with nuclear genomic structure, contact-zone dynamics, ecological differentiation, morphology and demographic history[76]. Evidence concerning assortative mating, hybrid fitness and barriers to effective gene flow would be particularly informative. We recommend retaining WH as the infraspecific taxon *P. b. polyctor* and treating the four ESUs as the principal evolutionary and conservation units pending such integrative assessment. Several peripheral lineages warrant particular attention, including WH (*polyctor*), YYM (*junia*), LNY and LYD (*kotoensis*), SWT (*stockleyi*) and SET (*pinratanai*) (Supplementary Table S1b).

Overall, *P. bianor* represents a spatially structured radiation in which historical admixture, regional isolation and peripheral differentiation have operated together. Its diversity is best understood as a hierarchy of lineages occupying different positions along the speciation continuum. Reproductive isolation can accumulate unevenly across ecological, genomic and mating-related dimensions[77], and the persistence of regional genomic structure alongside historical mixing makes this complex particularly informative for understanding how differentiated populations develop into independently evolving species.
